# Root-to-shoot ABA transport by NPF2.4 and NPF2.5 links stomatal regulation to photosynthetic performance

**DOI:** 10.64898/2026.10.07.757215

**Authors:** Wataru Yamori, Yuchen Qu, Alice Rossille, Christian Dubos, Yuri Kanno, Yumiko Takebayashi, Masami Y. Hirai, Shunsuke Watanabe, Mitsunori Seo

## Abstract

Fluctuating light strongly limits photosynthetic efficiency in plants, yet the mechanisms coordinating stomatal dynamics with rapid photosynthetic induction remain incompletely understood. Here, through a forward physiological screen in *Arabidopsis*, we identify the abscisic acid (ABA) transporters NPF2.4 and NPF2.5 as regulators of photosynthetic responses to fluctuating light. Loss-of-function *npf2.4* and *npf2.5* mutants exhibited accelerated stomatal opening and photosynthetic induction during light transitions, resulting in enhanced biomass accumulation and seed yield without compromising long-term drought tolerance. NPF2.4 and NPF2.5 mediated ABA uptake in yeast and were predominantly expressed in the root pericycle and cortex/epidermal tissues, respectively. Reciprocal grafting, xylem sap ABA measurements, and ABA treatment to the roots demonstrated that both transporters regulate root-to-shoot ABA transport. Our findings reveal that root-derived ABA modulates stomatal dynamics and photosynthetic efficiency under fluctuating light, establishing long-distance hormone transport as a key regulator of plant carbon gain in dynamic environments.

## Introduction

Abscisic acid (ABA) is a central plant hormone that coordinates plant growth and environmental responses by regulating processes, such as stomatal movement, photosynthesis, seed dormancy, and adaptation to abiotic stress^1,2^. Among these functions, the control of the stomatal aperture is particularly critical because it directly determines the balance between CO_2_ assimilation and transpirational water loss, thereby influencing carbon gain and plant productivity^3,4^. Although ABA-induced stomatal closure is classically associated with drought stress, accumulating evidence indicates that basal ABA signaling operates under non-stress conditions to fine-tune gas exchange and optimize photosynthetic performance^5,6^.

Endogenous ABA concentrations are tightly regulated by the balance between ABA biosynthesis and catabolism (inactivation). The rate-limiting step in ABA biosynthesis is catalyzed by 9-*cis*-epoxycarotenoid dioxygenase (NCED), whose expression responds dynamically to environmental cues^7^. Among several pathways for ABA catabolism reported to date, 8’-hydroxylation catalyzed by the CYP707A subfamily of P450 monooxygenases is the major and rate-limiting step^7^. However, metabolic regulation alone cannot account for the rapid and spatially restricted ABA responses observed in guard cells, particularly during dynamic environmental transitions, such as light fluctuations. This highlights the importance of transporter-mediated hormone redistribution.

According to the ion trap model, ABA, a weak acid, can move through the plasma membrane by passive diffusion^8,9^. However, the identification of dedicated transporters has demonstrated that ABA movement within plants is actively regulated^10,11^. ABA transport has long been discussed in the context of stomatal response to drought stress. Initial studies have reported the significance of root-derived ABA in the induction of stomatal closure in shoots (leaves) after partial root-zone drying^12,13^. However, later studies indicated that shoots (leaves), rather than roots, contribute largely to ABA accumulation and the regulation of stomatal aperture in leaves^14–17^. In leaves, vascular tissues, particularly phloem companion cells, appear to be the major sites of ABA biosynthesis, although guard cells can also synthesize ABA^18–22^. This indicates that ABA synthesized in vascular tissues is translocated to guard cells, thereby inducing stomatal closure. Several transporters involved in this process, such as the ATP-binding cassette (ABC) transporters ABCG25^23^ and ABCG40^24^, and NITRATE TRANSPORTER 1/PEPTIDE TRANSPORTER FAMILY (NPF) proteins NPF4.6^25^ and NPF5.1^26^ in Arabidopsis, have been identified. Despite these advances, the role of ABA transporters in the rapid stomatal responses to fluctuating light conditions remains largely unknown.

Here, through a forward genetic screen, we found that NPF2.4 and its close homolog NPF2.5, both of which exhibit cellular ABA uptake activities when expressed in yeast, play a role in photosynthetic regulation. Despite their effects on leaves, NPF2.4 and NPF2.5 were predominantly expressed in the root tissues. Integrated physiological analyses using grafting experiments and ABA measurements showed that NPF2.4 and NPF2.5 mediate root-to-shoot ABA transport, which modulates stomatal dynamics and enhances photosynthetic induction under fluctuating light without affecting steady-state photosynthetic capacity. These findings reveal a previously unrecognized mechanism by which root-derived ABA flux fine-tunes shoot gas exchange and optimizes plant growth.

## Results

### Identification of NPF2.4 through a photosynthesis-based screen

We performed a forward genetic screen focusing on physiological traits associated with the accelerated activation of photosynthetic electron transport upon illumination to identify novel regulators of photosynthetic performance (Fig. S1). Mutants displaying faster photosynthetic induction than the wild-type were selected for further analysis. Through this screening, we isolated a mutant line exhibiting enhanced photosynthetic induction responses compared with wild-type plants from a pool of T-DNA insertion lines obtained from the Arabidopsis Biological Resource Center (ABRC).

Genomic sequences flanking the T-DNA insertion were amplified using thermal asymmetric interlaced PCR (TAIL-PCR) to determine the gene responsible for this phenotype. Sequence analysis revealed that the T-DNA insertion occurred within the *NPF2.4* gene (AT3G45700) (Fig. S2), and the mutant was designated as *npf2.4-1*. We obtained an independent T-DNA insertion line for *NPF2.4* [*npf2.4-2* (SALK_005261)] to confirm this phenotype. Because *NPF2.4* shares high sequence similarity with the closely related transporter *NPF2.5* (AT3G45710), we obtained two independent T-DNA insertion mutants for *NPF2.5* [*npf2.5-1* (GK-936H04) *and npf2.5-2* (SM_3_31001)] to evaluate their roles in regulating stomatal behavior and photosynthetic performance.

### Accelerated dynamic photosynthesis under fluctuating light

We analyzed the gas-exchange responses of *npf2.4* and *npf2.5* loss-of-function mutants under several independent fluctuating light regimes to investigate whether NPF2.4 and NPF2.5 are involved in dynamic photosynthetic regulation. Under all three independent light transition patterns tested (Fig. 1A, E, I), both mutants showed a more rapid induction of stomatal conductance and CO_2_ assimilation following transitions from low to high light (Fig. 1B, C, F, G, J, K). Elevated intercellular CO_2_ concentration (Ci) in the mutants further indicated improved CO_2_ diffusion into leaves (Fig. 1D, H, L). Importantly, the steady-state light-response curves did not differ between the two genotypes (Fig. S3), indicating that NPF2.4 and NPF2.5, rather than maximum photosynthetic capacity, specifically affected dynamic photosynthetic responses. Chlorophyll fluorescence imaging results corroborated these findings (Fig. S4). Under fluctuating light conditions, *npf2.4* and *npf2.5* displayed higher effective quantum yields of PSII [Y(II)] and faster induction kinetics (Fig. S4). These data demonstrated that NPF2.4 and NPF2.5 contribute to efficient photosynthetic induction during light transitions.

**Figure 1.**
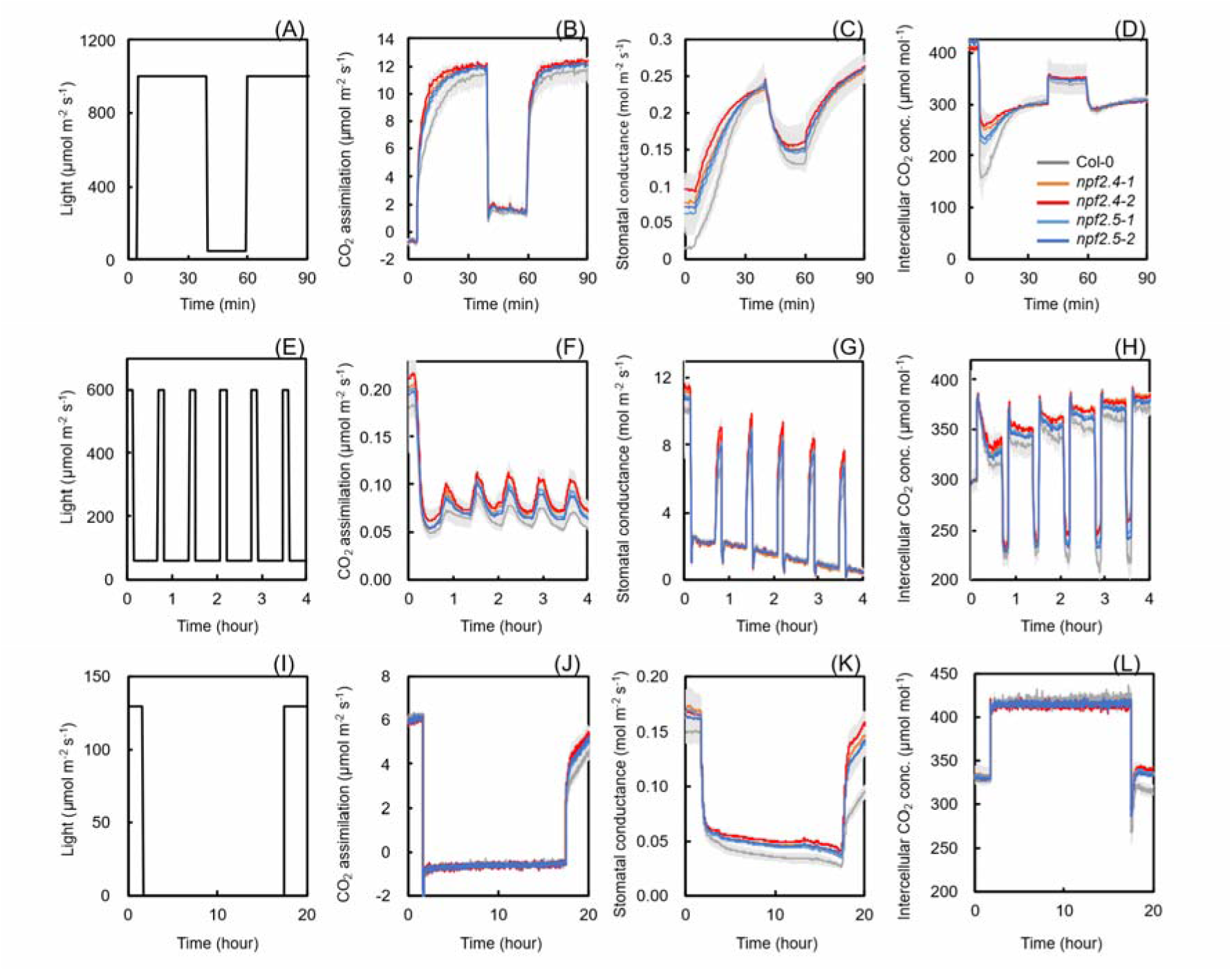
Photosynthetic responses to fluctuating light. (A) Light fluctuation regime used for gas-exchange measurements. (B) CO_2_ assimilation rate under the light fluctuation shown in (A). (C) Stomatal conductance under the light fluctuation shown in (A). (D) Intercellular CO_2_ concentration under the light fluctuation shown in (A). (E) Light fluctuation regime used for gas-exchange measurements. (F) CO_2_ assimilation rate under the light fluctuation shown in (E). (G) Stomatal conductance under the light fluctuation shown in (E). (H) Intercellular CO_2_ concentration under the light fluctuation shown in (E). (I) Light fluctuation regime used for gas-exchange measurements. (J) CO_2_ assimilation rate under the light fluctuation shown in (I). (K) Stomatal conductance under the light fluctuation shown in (I). (L) Intercellular CO_2_ concentration under the light fluctuation shown in (I). Values represent mean ± SEM. (*n* = 6).

### Enhanced stomatal opening and growth in *npf2.4* and *npf2.5* mutants

We analyzed the leaf surface temperature and growth phenotypes to assess whether altered dynamic photosynthesis translated into changes in stomatal regulation and plant performance. Thermal imaging of 25-day-old plants revealed that all *npf2.4* and *npf2.5*-mutant lines exhibited significantly lower leaf surface temperatures than wild-type Col-0 (Fig. 2A, B; Fig. S5). As reduced leaf surface temperature reflects enhanced evaporative cooling, these results suggest increased stomatal opening in the mutants. Increased stomatal conductance is accompanied by increased growth. Both shoot and root biomass were significantly higher in *npf2.4* and *npf2.5* mutants than in Col-0 40 days after sowing (Fig. 2C–E). This growth advantage persisted through the reproductive stage, during which mutants exhibited increased shoot biomass and higher seed yield (Fig. 2F–H). These findings indicated that the loss of NPF2.4 or NPF2.5 functions promotes vegetative and reproductive growth under non-stress conditions.

**Figure 2.**
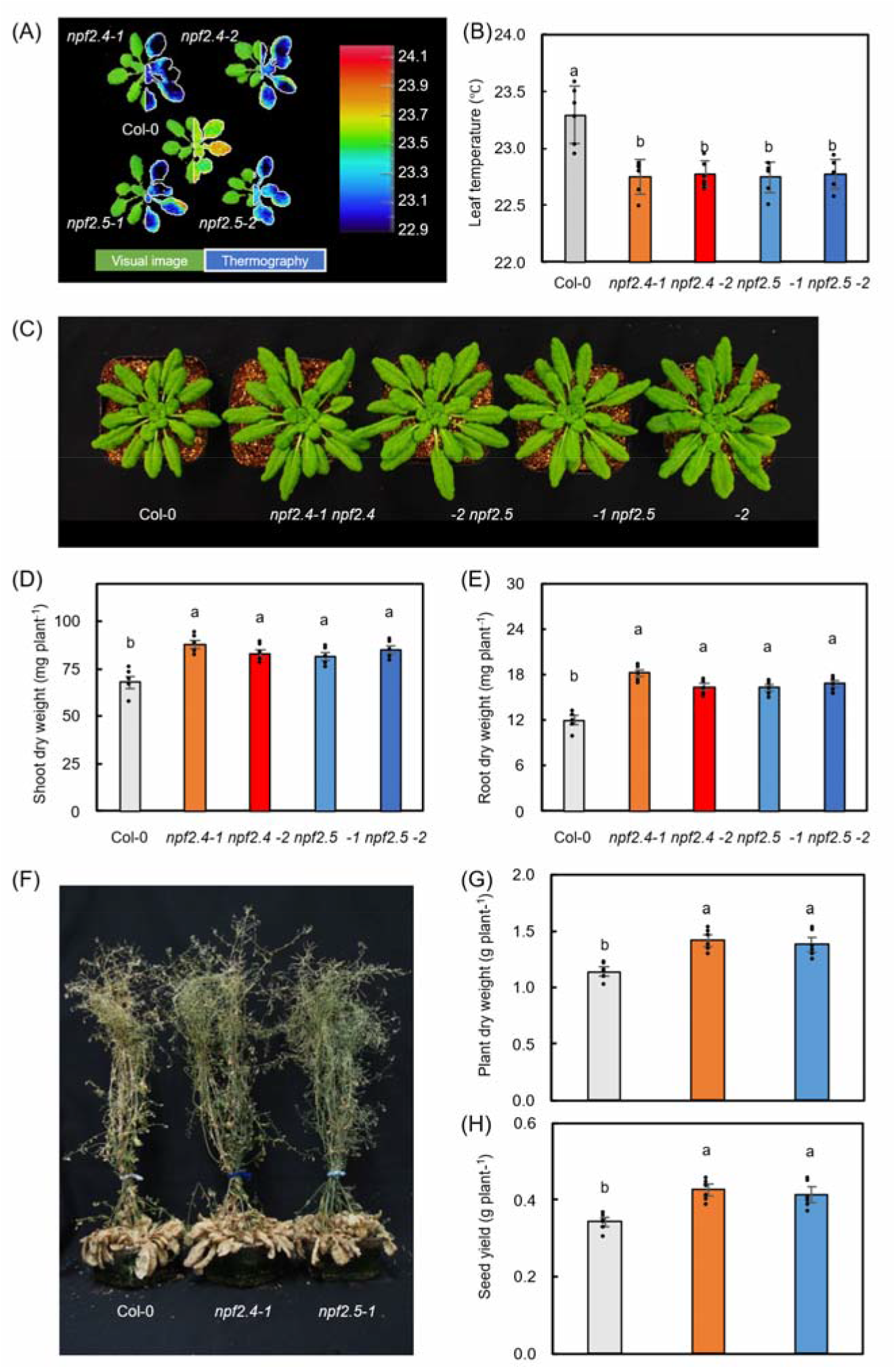
Plant growth, thermal imaging, and biomass accumulation. (A) Representative images of plant growth (left) and corresponding thermal images obtained using infrared thermography (right) of 25-day-old plants. Thermal images were acquired 1 h after the onset of illumination under growth-light conditions. The original thermal imaging data are shown in Fig. S3. (B) Leaf surface temperatures quantified from thermal images shown in (A), measured using a far-infrared camera. (C) Representative images of 40-day-old plants. (D) Dry weight of above-ground parts at 40 days after sowing. (E) Dry weight of below-ground parts at 40 days after sowing. (F) Representative images of 2.5-month-old plants. (G) Dry weight of above-ground parts at 2.5 months after sowing. (H) Seed yield per plant at 2.5 months after sowing. Values represent mean ± SEM. Different letters indicate significant differences at *P* < 0.05, as determined using the Tukey–Kramer HSD test (*n* = 6).

### NPF2.4 and NPF2.5 mediate cellular ABA uptake

Given that several NPF proteins function as ABA transporters^11^ and that stomatal phenotypes have been observed in mutants defective in ABA transport^2^, we hypothesized that NPF2.4 and NPF2.5 transport ABA. Using a modified yeast two-hybrid (Y2H) system with the ABA receptor complex that we established previously^25^, we compared the potential ABA transport activities of 45 of the 53 Arabidopsis NPF proteins expressed in yeast cells based on their growth in liquid selection medium containing a low concentration (50 nM) of ABA (Fig. S6). Consistent with its initial identification as a protein capable of inducing the interaction between the ABA receptor PYR and PP2C protein phosphatase in this Y2H system, the ABA transporter NPF4.6 was the most efficient in promoting yeast growth. In addition, close homologs of NPF4.6, as well as some members of the NPF5 clade, including the ABA transporter NPF5.1, promoted yeast growth in this assay system. Furthermore, NPF2.4, NPF2.5, and other NPFs in clade 2 exhibited potential ABA transport activities, albeit to a lesser extent. These results indicate that NPF2.4 and NPF2.5, among the NPFs tested, have relatively high affinity for ABA. Finally, we confirmed that NPF2.4 and NPF2.5 indeed mediate cellular ABA uptake by quantifying ABA uptake in yeast cells using mass spectrometry (Fig. 3).

**Figure 3.**
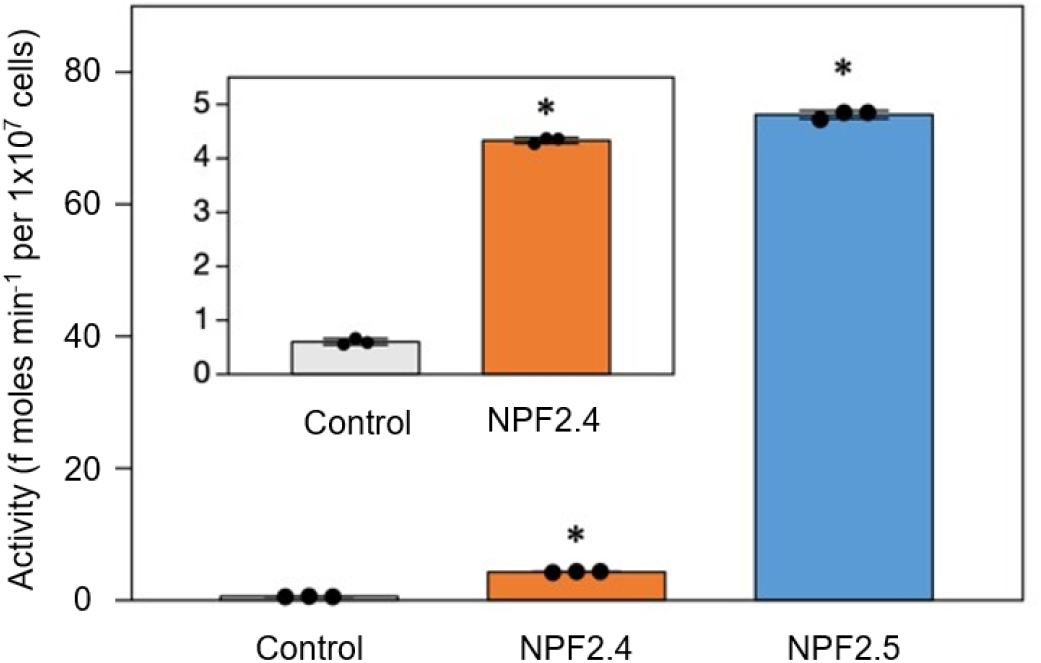
ABA transport activities of Arabidopsis NPFs determined using the direct assay. Yeast cells expressing NPF2.4 or NPF2.5 were incubated with solutions containing 10 μM ABA, and ABA taken up by the cells was quantified by using LC-MS/MS. Yeast cells transformed with an empty vector were used as controls. Values represent mean ± SD. Asterisks indicate significant differences at *P* < 0.05, as determined using Dunnett’s multiple comparison test (*n* = 3).

### Localization of NPF2.4 and NPF2.5 in roots

To investigate the spatial expression patterns of *NPF2.4* and *NPF2.5*, we generated transgenic plants expressing GUS under the control of the *NPF2.4* or *NPF2.5* promoter (*pNPF2.4:GUS* and *pNPF2.5:GUS*). In *pNPF2.4:GUS* plants, GUS staining was observed in the distal region of the primary root, whereas no staining was detected at the root tip (Fig. 4A). Lateral roots were also stained; however, their tips showed no detectable GUS signals. The inner tissues around the vasculature were stained in both the primary and lateral roots. In *pNPF2.5:GUS* plants, GUS staining was observed throughout most regions of the primary root; however, little or no staining was detected in the tip regions (Fig. 4B). The basal regions of the lateral roots were also stained, but GUS signals were not detected in their tip regions. In contrast to *pNPF2.4:GUS*, the outer tissues surrounding the vasculature were sained in both primary and lateral roots. Transgenic lines expressing GFP-tagged NPF2.4 or NPF2.5, driven by their respective promoters (*pNPF2.4:gDNA-GFP* and *pNPF2.5:gDNA-GFP*) were generated to more precisely determine where NPF2.4 and NPF2.5 function. For these GFP-tagged constructs, genomic DNA was inserted into a binary vector instead of cDNA. In *pNPF2.4:gDNA-GFP* plants, GFP signals were observed in the pericycle cells (Fig. 5A). In contrast, GFP signals were detected in the cortex and, to a lesser extent, in the epidermal cells of *pNPF2.5:gDNA-GFP* plants (Fig. 5B). These protein localization patterns were consistent with those predicted using GUS staining. In both *pNPF2.4:gDNA-GFP* and *pNPF2.5:gDNA-GFP* transgenic lines, GFP signals were detected at the plasma membrane with no apparent polarity (Fig. 5; Fig. S7).

**Figure 4.**
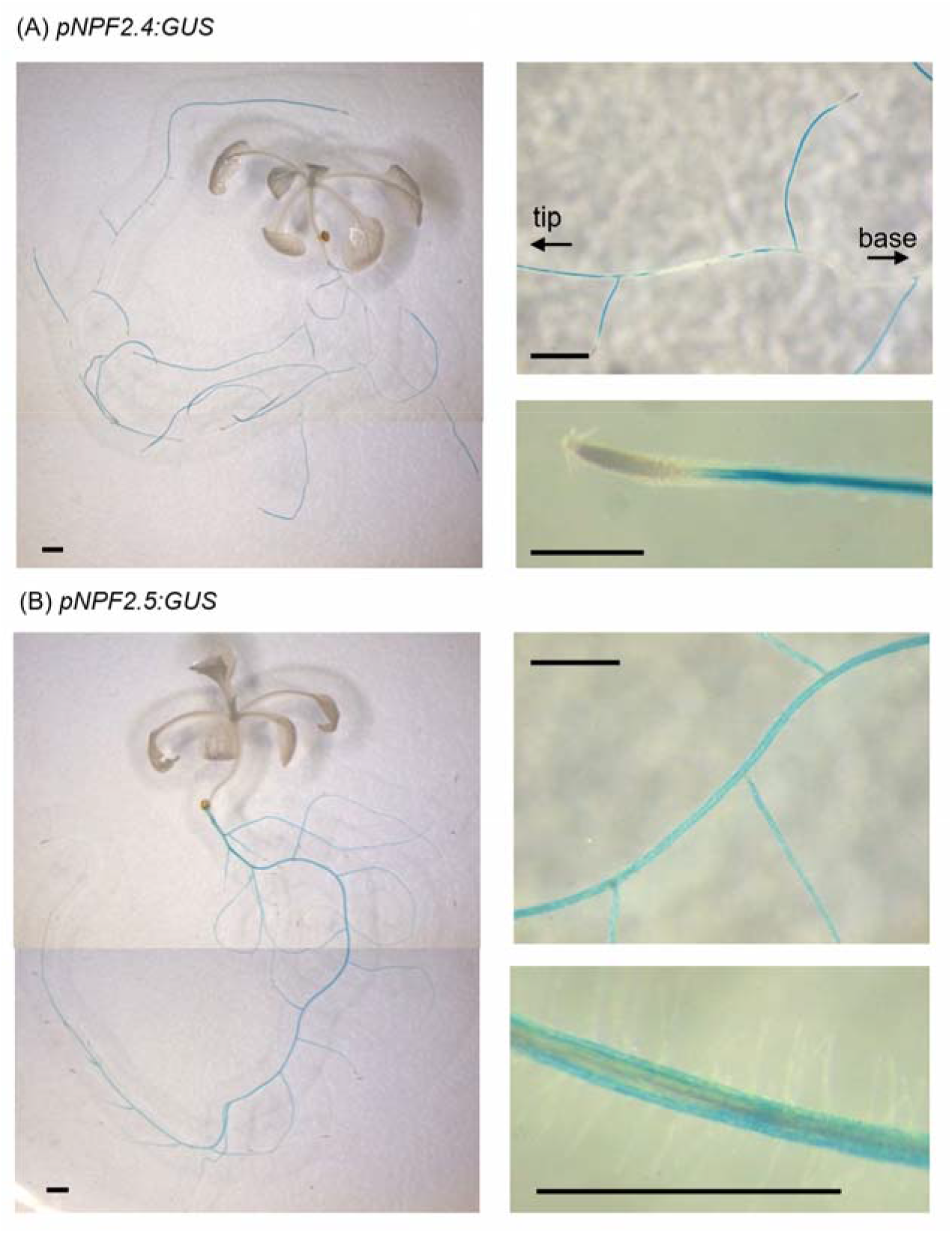
Spatial expression patterns of *NPF2.4* and *NPF2.5*. (A) GUS staining of *pNPF2.4:GUS* plants. (B) GUS staining of *pNPF2.5:GUS* plants. Arrows in (A) indicate the direction of the main root toward the root tip (tip) or basal (base) regions. Bars indicate 1 mm. GUS staining was performed on 2-week-old plants.

**Figure 5.**
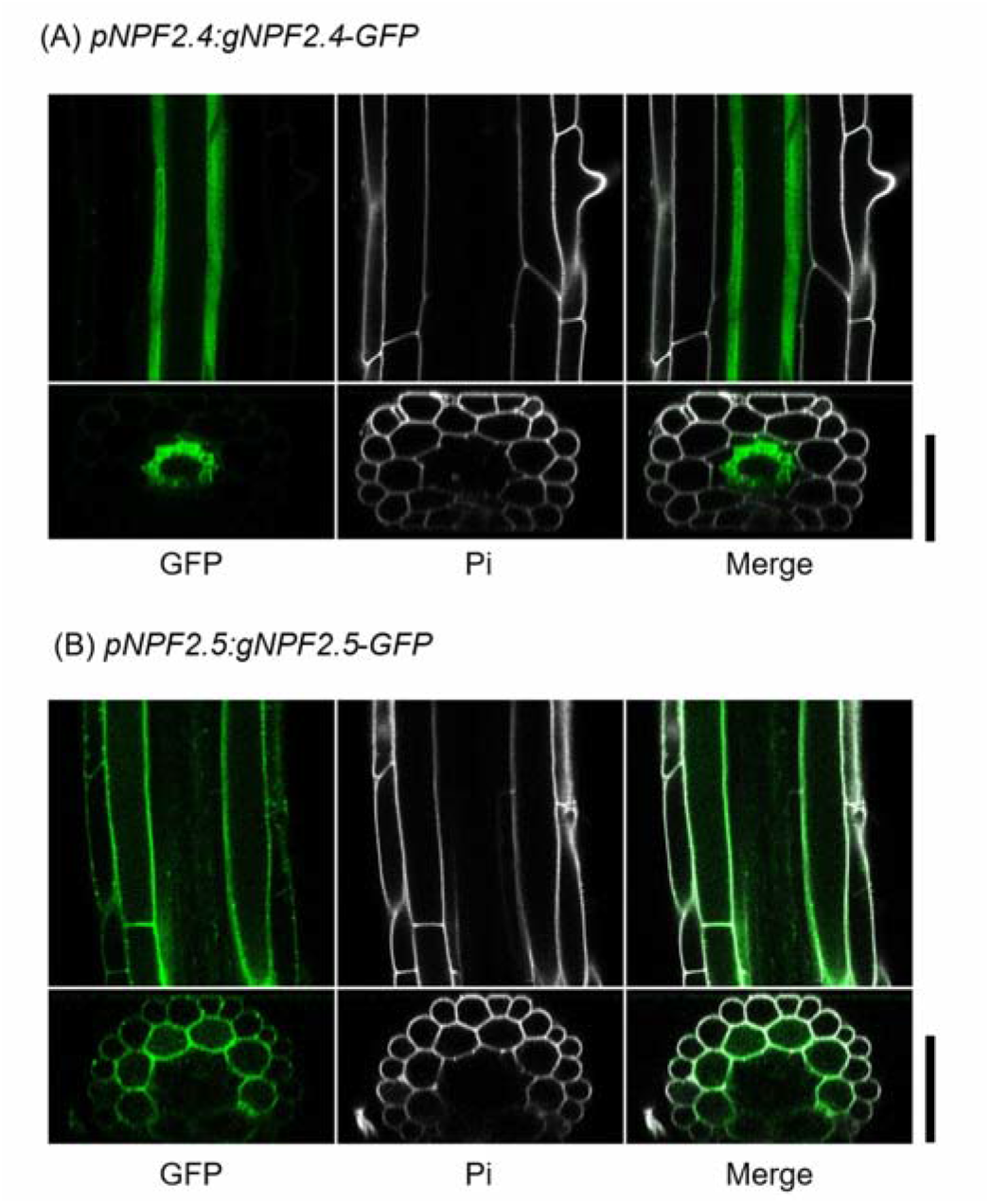
Localization of NPF2.4 and NPF2.5 proteins in root tissues. (A) GFP signals detected in roots of 10-day-old *pNPF2.4:gDNA-GFP* plants. (B) GFP signals detected in roots of 10-day-old *pNPF2.5:gDNA-GFP* plants. Pi shows the staining of the cell walls with propidium iodide. The upper and lower panels show the longitudinal and transverse sections, respectively. Bars indicate 50 μm.

### NPF2.4 and NPF2.5 expressed in roots determine shoot growth

We performed reciprocal grafting experiments between wild type and *npf2.4* and between wild type and *npf2.5* to determine whether the expression of NPF2.4 and NPF2.5 detected in roots was responsible for shoot physiology. Consistent with the results presented in Fig. 2, self-grafted *npf2.4* and *npf2.5* plants exhibited greater shoot biomass than the self-grafted wild-type (Fig. 6). However, plants grafted with *npf2.4* or *npf2.5* rootstocks, but not with the corresponding scions, grew larger than the self-grafted wild type (Fig. 6). These data indicate that the increased biomass associated with enhanced photosynthesis observed in *npf2.4* and *npf2.5* results from defects in NPF2.4 and NPF2.5, respectively.

**Figure 6.**
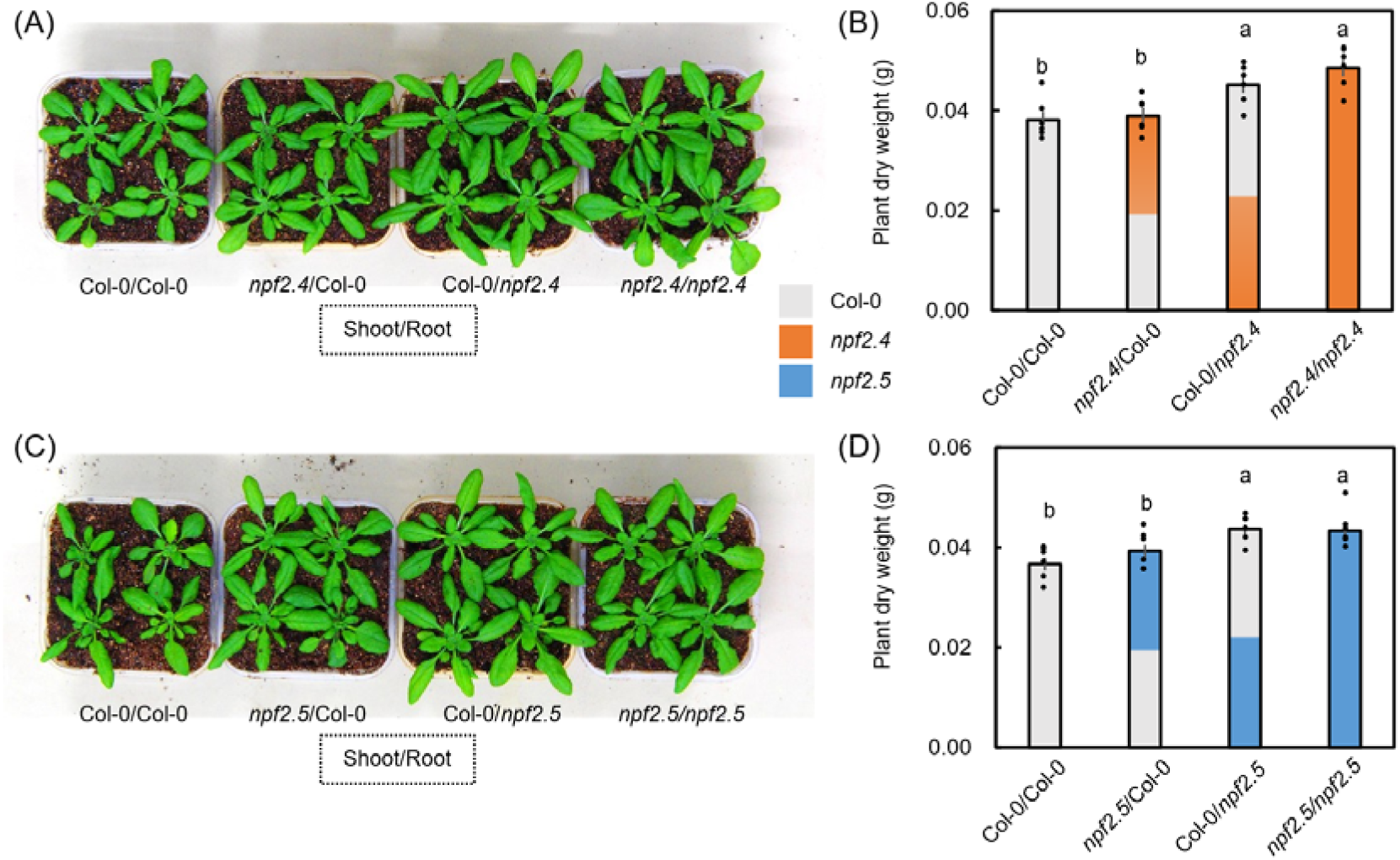
Growth performance of grafted plants with npf2.4 or npf2.5 in the scion or rootstock. (A) Representative images of grafted plants combining Col-0 and npf2.4 genotypes. (B) Shoot dry weight of grafted plants shown in (A), measured 35 days after transplanting. (C) Representative images of grafted plants combining Col-0 and npf2.5 genotypes. (D) Shoot dry weight of grafted plants shown in (C), measured 35 days after transplanting. Seedlings were grafted 10 days after sowing, transplanted 18 days after sowing, photographed, and harvested 48 days after sowing. Values represent mean ± SEM. Different letters indicate significant differences at *P* < 0.05, as determined using the Tukey–Kramer HSD test (*n* = 6).

Given that NPF2.4 and NPF2.5 function in the roots and mediate transmembrane ABA transport, we hypothesized that they may be involved in long-distance ABA transport from the roots to shoots. To test this possibility, we quantified the ABA levels in the xylem sap. Under well-watered conditions, xylem sap ABA levels were comparable among the wild type, *npf2.4,* and *npf2.5*. However, when the plants were exposed to mild drought stress by withholding water for 5 d, a significant increase in xylem sap ABA concentration was observed in the wild type, but not in *npf2.4* or *npf2.5* (Fig. 7). We then examined the effects of exogenous ABA applied to the roots on the leaf surface temperature, which is associated with stomatal aperture. While ABA application to the roots significantly increased leaf surface temperature in the wild type, the treatment did not affect leaf surface temperature in *npf2.4* and *npf2.5* (Fig. 8). More importantly, such changes are not caused by changes in stomatal size nor density, as microscope determinations showed no significant differences in these two parameters (Fig. S8). These data support the hypothesis that *npf2.4* and *npf2.5* are defective in root-to-shoot ABA transport.

**Figure 7.**
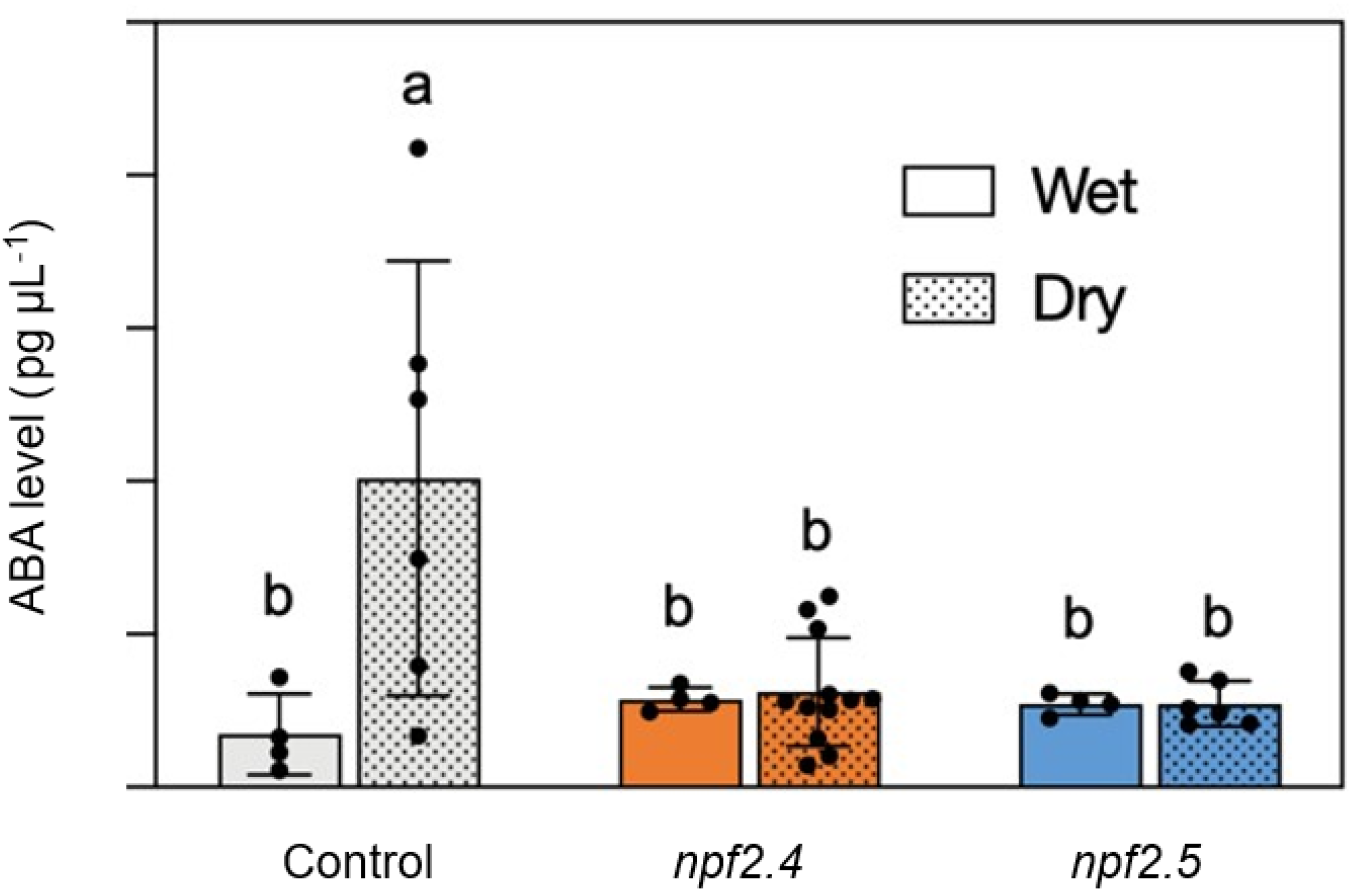
ABA levels in xylem sap of wild type, *npf2.4-1* and *npf2.5-1*. Xylem sap was extracted from 40-day-old rosette leaves of wild-type (WT), *npf2.4-1* (*npf2.4*), and *npf2.5-1* (*npf2.5*) plants before (Wet) or after (Dry) drought stress and subjected to quantification of ABA by LC-MS/MS. The bars indicate standard deviations. Values represent mean ± SD. Different letters indicate significant differences at *P* < 0.05, as determined using the Tukey–Kramer HSD test [n = 4 (WT, Wet), 6 (WT, Dry), 4 (*npf2.4*, Wet), 12 (*npf2.4*, Dry), 4 (npf2.5, Wet), and 6 (npf2.5, Dry)].

**Figure 8.**
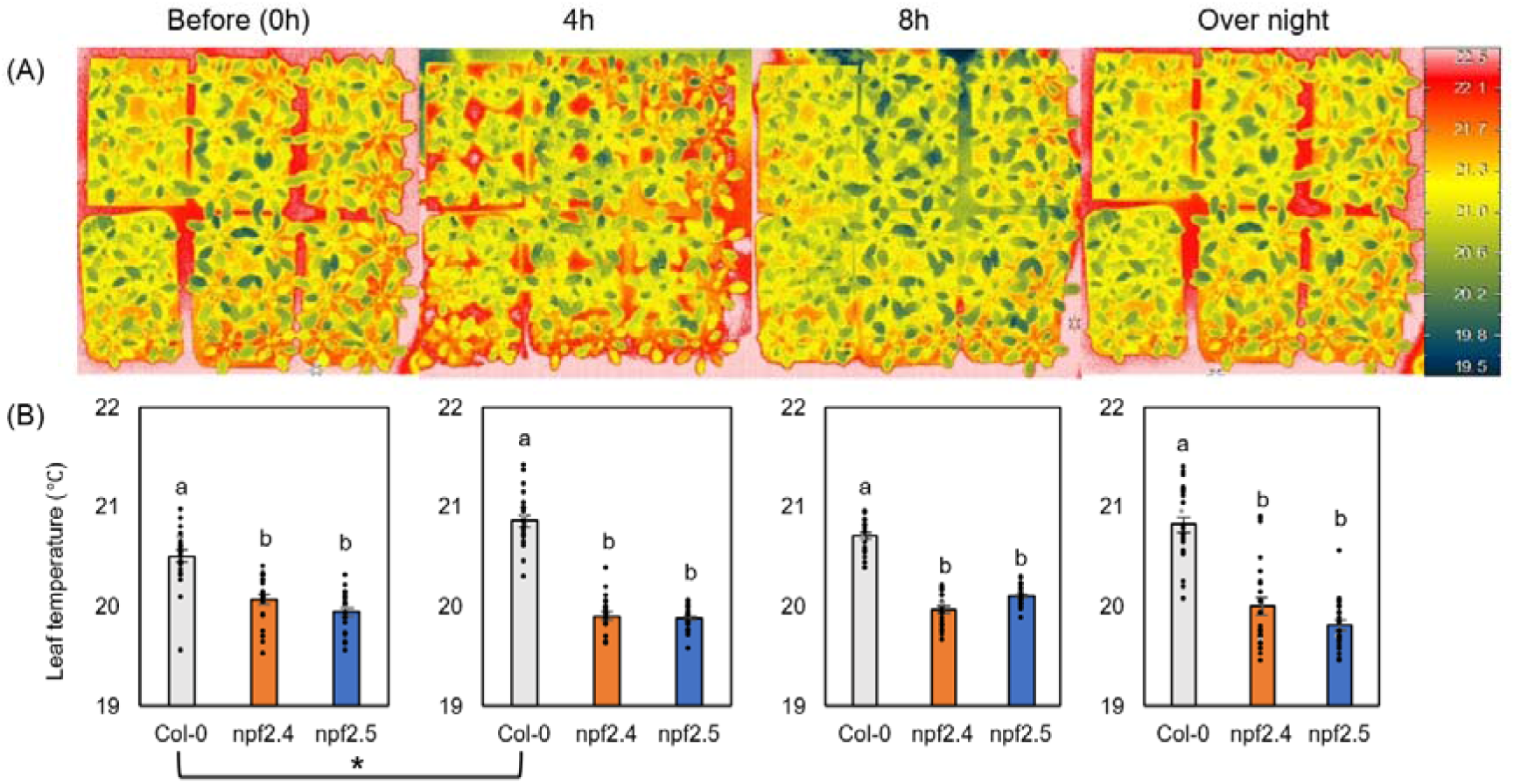
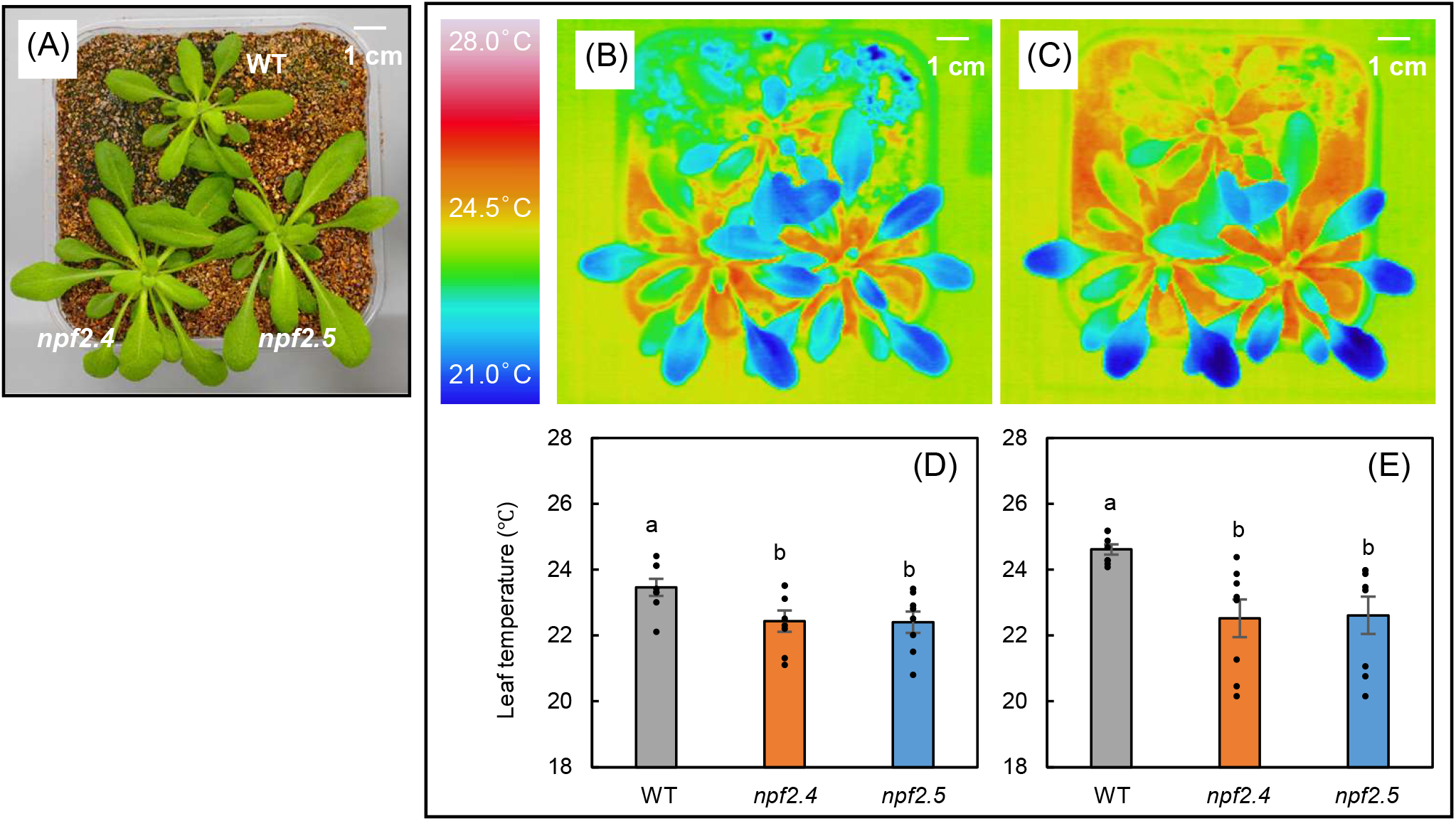
Changes in leaf temperature in Arabidopsis thaliana Col-0, npf2.4, and npf2.5 plants following root application of abscisic acid (ABA). (A) Representative thermal images of Col-0, npf2.4, and npf2.5 plants before ABA treatment (0 h) and at 4 h, 8 h, and overnight after root application of ABA. Leaf temperature was monitored using an infrared thermal imaging camera under the same experimental conditions. (B) Quantification of leaf temperature at the indicated time points before and after ABA treatment. Statistical significance was assessed by two-way ANOVA followed by Tukey’s multiple-comparison test (P < 0.05). Different lowercase letters indicate significant differences among means, n=24. Asterisks indicate significant differences at *P* < 0.05

Some members of NPF proteins have been reported to transport nitrate (NO□□)^27^. In addition, NPF2.4 functions as a chloride (Cl□) transporter^28, 29^. Therefore, it might be possible that the phenotypes observed in *npf2.4* and *npf2.5* were caused by their defects in NO□□ and/or Cl□ transport. However, no significant differences in NO□□ and Cl□ contents were detected between wild type and the *npf2.4* or *npf2.5* mutants (Fig. S9A, B). In addition, total leaf nitrogen content, RuBisCO protein abundance, and chlorophyll levels were comparable among the genotypes (Fig. S9C-E). These results indicated that the enhanced photosynthetic performance and growth observed in the mutants were unlikely to result from altered nutrient status or photosynthetic capacity, but rather from changes in ABA transport and stomatal regulation.

## Discussion

In this study, we found that mutants defective in NPF2.4 and its close homolog NPF2.5 (*npf2.4* and *npf2.5*, respectively) exhibited moderately increased stomatal aperture and enhanced photosynthetic rates during the transition from dark to light (photosynthetic induction), which is reminiscent of ABA transporter mutants such as *abcg25* and *abcg40*^2^. In addition, several members of the NPF protein family are involved in ABA transport^25,26,30^. These results prompted us to examine whether NPF2.4 and NPF2.5 function as ABA transporters. Y2H-based indirect assays indicated that NPF2.4 and NPF2.5, among the 45 Arabidopsis NPFs tested, exhibited relatively high affinity or specific activity for ABA (Fig. S6). Notably, NPF4.6, NPF5.1, and NPF2.13, whose functions as ABA transporters have been demonstrated experimentally, efficiently promoted yeast growth in the same assay. Further direct transport using with mass-spectrometry confirmed that both NPF2.4 and NPF2.5 mediate cellular ABA uptake (Fig. 3).

Earlier studies have indicated that leaf stomatal closure is induced by partial root-zone drying, and that ABA is the root-derived signal involved in this process^12,13^. However, recent studies have shown that ABA synthesized in shoots (leaves), rather than in roots, plays a predominant role in stomatal regulation^16,17^. In Arabidopsis, ABA biosynthesis enzymes such as NCED3, ABA2, and AAO3 are preferentially expressed in leaf vascular tissues (most likely in phloem companion cells)^18,19,20^, indicating that the hormone is actively synthesized in this tissue type. This indicates that the ABA synthesized in vascular tissues must be translocated to the guard cells to close the stomata. Several transporters involved in this process have been identified. For example, in Arabidopsis, ABCG25 mediates ABA export from its biosynthesis site (vascular tissues), whereas ABCG40 and NPF4.6 facilitate ABA uptake into guard cells^23,24,26^. Given that *npf2.4* and *npf2.5* mutants show altered stomatal apertures, NPF2.4 and NPF2.5 may be involved in ABA transport from vascular tissues to guard cells. Nevertheless, our promoter-reporter analysis showed that the two transporters were mainly expressed in the root tissues (Fig. 4 and 5). Reciprocal grafting confirmed that NPF2.4 and NPF2.5 expression in roots is required for proper stomatal regulation in the shoots (Fig. 6). NPFs are a nitrate transporter family^31^. Additionally, NPF2.4 and NPF2.5 have been characterized as chloride transporters^28,29^. However, endogenous levels of nitrate and chloride ions, as well as total nitrogen and chlorine content in leaves, were similar among the wild type, *npf2.4*, and *npf2.5* (Fig. S9). In contrast, quantification of ABA in the xylem sap, together with investigation of stomatal responses to ABA applied to roots (Fig. 7 and 8), suggests that NPF2.4 and NPF2.5 are involved in root-to-shoot translocation of ABA. These findings led us to reconsider the significance of root-derived ABA in stomatal regulation, even though it plays a minor role compared to the ABA synthesized in leaf vascular tissues. Notably, the identification of NPF2.4 and NPF2.5 as novel ABA transporters, originated from a screen based on photosynthesis-related phenotypes rather than from classical screens based on hormone- or stress-related phenotypes, highlighting the power of this approach to identify previously unrecognized regulatory mechanisms.

To date, several root-derived signaling molecules that regulate stomatal apertures, such as CLE peptides^32^ and sulfate^33^, have been identified. It is important to investigate how these molecules integrate multiple signals and coordinately regulate stomatal aperture in a changing environment. Importantly, the enhanced photosynthetic performance observed in *npf2.4* and *npf2.5* mutants was not associated with changes in stomatal density or size, nor in the steady-state photosynthetic capacity, RuBisCO abundance, or chlorophyll content (Fig. S8, S9), indicating that the mutants did not affect basal biochemical capacity. Notably, the phenotypes were observed specifically under fluctuating light conditions, particularly during photosynthetic induction, without affecting drought stress resistance (Fig. S10). This contrasts with typical mutants affected by ABA biosynthesis or signaling, which exhibit an increased stomatal aperture but reduced stress tolerance. These observations suggest that NPF2.4/NPF2.5-mediated ABA transport specifically influences dynamic stomatal regulation under non-stress conditions. This mechanism may represent an adaptive strategy by which plants optimize productivity through the fine control of long-distance hormone flux while retaining the ability to mount rapid stress responses.

ABA is often regarded as a stress hormone; however, accumulating evidence indicates that basal ABA levels are essential for optimizing stomatal behavior and photosynthetic efficiency^5,6^. Subtle modulation of ABA availability in guard cells can shift the balance between CO_2_ uptake and water loss, thereby influencing carbon gain, even in the absence of environmental stress^4^. Our results suggest that NPF2.4- and NPF2.5-mediated ABA transport from the roots helps maintain this balance. The reason why two different sources of ABA, one synthesized in the leaf vascular tissues and the other derived from the roots, are required for stomatal regulation remains unclear. However, ABA from different sources may have distinct mobility within plant tissues. ABA synthesized in vascular tissues must be exported from the cells at least once after its synthesis. In contrast, root-derived ABA can potentially move through the apoplastic space with the transpiration stream to reach guard cells.

Although the involvement of NPF2.4 and NPF2.5 in root-to-shoot ABA transport is evident, the mechanisms by which these transporters regulate this process remain unclear. The expression patterns of ABA biosynthesis genes and enzymes suggest that ABA is synthesized in the vascular tissues of roots and leaves^18,20^. It has also been reported that ABA synthesized in leaves is translocated to the roots via the phloem^16,17,34^. Based on these observations, we speculated that NPF2.4 functions to retain ABA synthesized in and translocate to root vascular tissues within pericycle cells, enabling ABA concentrated in this cell type to be loaded into the xylem for long-distance root-to-shoot transport, which also involves the ABA exporter ABCG25^35^ (Fig. S11). Unexpectedly, despite the similar stomatal phenotypes observed in *npf2.4* and *npf2.5*, the expression pattern of NPF2.5 was different from that of NPF2.4 (Fig. 4 and 5). It appears that NPF2.5 in epidermal/cortex cells is not directly related to ABA loading into the xylem. ABA transport from the inner to outer tissues within the roots has also been reported^36^. In contrast, ABA exogenously applied to roots can be transported to the shoots to induce stomatal closure (Fig. 7 and 8). Therefore, ABA is likely transported radially in both inward and outward directions in the roots. In this model, NPF2.5 might be associated with other ABA transporters to coordinately regulate radial ABA transport, which might be coupled with ABA loading into the xylem (Fig. S11).

Although our data established that NPF2.4 and NPF2.5 are key regulators of root-to-shoot ABA transport and, hence, photosynthetic induction, several questions remain to be addressed. Public databases indicate that the transcript levels of *NPF2.4* and *NPF2.5* remain relatively constant across different photoperiods and under abiotic stress conditions.^37^ (Fig. S12). However, the post-transcriptional/translational regulation of NPF2.4 and NPF2.5, which may affect its transport activity, has not been examined. Examination of the substrate specificity of NPF2.4 and NPF2.5, particularly for ABA precursors, will also be important for understanding their in vivo functions. Finally, determining whether analogous NPF-mediated ABA transport mechanisms operate in crop species could provide new strategies to enhance dynamic photosynthetic performance and yield stability under field conditions by targeting hormone transport pathways.

## Materials and Methods

### Plant Materials and Growth Conditions

*Arabidopsis thaliana* wild-type (WT, Col-0) and two independent T-DNA insertion mutants for NPF2.4, *npf2.4-1* (isolated by in-house screening), and *npf2.4-2* (SALK_005261C), as well as two independent NPF2.5 mutants, *npf2.5-1* (GK-936H04) and *npf2.5-2* (SM_3_31001), were used in this study (Fig. S2). The plants were cultivated individually in plastic pots (280 or 380 mL) containing a 1:1 (v/v) mixture of vermiculite and commercial nutrient soil (Metromix 350; HYPONeX, Japan). Growth was conducted under controlled environmental conditions at 23 °C with 65% relative humidity, a 10-h light/14-h dark photoperiod, and a photosynthetic photon flux density of 130 μmol m^−2^ s^−1^, under ambient CO_2_ concentration (approximately 400 μmol mol^−1^). The pots were arranged randomly on plastic trays, and their positions were periodically rotated to minimize positional effects. The plants were irrigated once weekly to prevent desiccation.

For the generation of *pNPF2.4:GUS* plants, a promoter region (2 kb upstream of the ATG start codon) of *NPF2.4* was amplified by PCR with primer pair 5□-GGGGACAAGTTTGTACAAAAAAGCAGGCTTCTTTGGACCACTTGAGACC-3□/5□-GGGGACCACTTTGTACAAGAAAGCTGGGTGGCAGAGAACAAGTGGAAGC-3□ and cloned into pDONR207. After confirming the sequence, the promoter region was cloned into the pGWB3^38^. For the generation of *pNPF2.5:GUS* plants, a promoter region (2 kb upstream of the ATG start codon) of NPF2.4 was amplified by PCR with primer pair 5□-GCCGCCCCCTTCACCTTCCTGCTCTCGTGATTAATGGAATAGGTG-3□/5□-GGCGCGCCCACCCTTGGTAGAGAACAAGATGAACCAGGAGGGCAA-3□ and cloned into p-ENTR-D-TOPO. After confirming the sequence, the promoter region was cloned into pGWB3. Transgenic plants carrying these constructs were obtained via Agrobacterium-mediated transformation using the floral dipping method.

For te generation of *pNPF2.4:gDNA-GFP* and *pNPF2.5:gDNA-GFP* plants, the 2.0 kb promoter region and the whole 2.3 kb genomic region of NPF2.4, and the 1.3 kb promoter region and the whole 2.0 kb genomic region of NPF2.5, were amplified by PCR using primer pairs 5□-AAAAGCAGGCTCCACCATGAAAGCTTACTAGTTGTTAAACGTTTCC-3□/5□-CTGGGTCTAGATATCTCGAAGTTTGTAACATCTTTAAGATCTTGT-3□ and 5□-AAAAGCAGGCTCCACCATGCTTTTTTAAGGATTCTTCACCTCCT-3□/5□-CTGGGTCTAGATATCTCGAAGGTTTTAACATCTTTAGGATCTTGT-3, respectively. The amplicons were cloned into pENTR4 using Gibson assembly. After sequence confirmation, the *pNPF2.4:gDNA* and *pNPF2.5:gDNA* sequences in the pENTR4 vector were transferred into pFAST-R07^39^. Transgenic plants carrying these constructs were generated via Agrobacterium-mediated transformation using the floral dip method.

### Gas Exchange Measurements

Gas exchange measurements were performed to evaluate dynamic stomatal regulation and photosynthetic induction responses. Photosynthetic induction analysis was conducted overnight in the dark-adapted plants. Fully expanded young leaves from 6–8-week-old plants were selected for the measurements. Gas exchange parameters were recorded at 30-s intervals using an open gas exchange system (LI-6400XT; Li-COR, USA). Net CO_2_ assimilation rate (A), stomatal conductance to water vapor (gs), and intercellular CO_2_ concentration (Ci) were simultaneously monitored to assess the coordination between stomatal opening and CO_2_ diffusion during the transition from dark or low-light to high-light conditions. Measurements were conducted under controlled conditions with a block temperature of 25 °C, relative humidity maintained between 55% and 65%, a reference CO_2_ concentration of 400 μmol mol^−1^, and an airflow rate of 500 μmol s^−1^. Leaves were initially maintained in darkness for 5 min and subsequently exposed to a photosynthetic photon flux density (PPFD) of 1000 μmol m^−2^ s^−1^ for 40 min to induce photosynthesis.

### Stomatal apertures parameters

Leaf surface temperatures were measured in 5-6-week-old plants using a far-infrared thermal camera (Thermo Tracer TH9100MR; NEC Avio Infrared Technologies, Japan). Leaf temperature was used as a proxy for transpirational cooling associated with stomatal apertures. For root zone related ABA treatment, 3-4-week-old plants were cultivated in cell tray (volume each cell), and a dose of 1 mL 10 μM (±)-ABA solution (A1049; Sigma-Aldrich Co. LLC, USA) was injected into the soil of each cell, allowing uptake through the root system. Thermal images were acquired before ABA injection, 4 hours, 8 hours after ABA injection, and on the following day (overnight).

Stomatal density was measure using 5-6-week-old plants. Transparent nail varnish was applied on fully expanded Arabidopsis leaf upper surface to acquire the cast of the surface structure of determined locations. Solidified nail varnish was then carefully removed from the leaf and recorded through a 200x microscope, then density was determined by counting the number of stomata within each field of view.

Stomatal size was measured using the nail varnish cast acquired above. A SEM () was used to record the 3D structure of stomata and the size was determined by the length of the stomatal pore along its longitudinal axis.

### Chlorophyll Fluorescence Measurements

The chlorophyll fluorescence parameters were measured using a pulse amplitude-modulated fluorometer (IMAG-MAX/L; Heinz Walz GmbH, Germany). The maximum quantum efficiency of PSII (Fv/Fm) was used to assess the intrinsic photochemical capacity of PSII after adaptation to dark conditions. The effective quantum yield of PSII [Y(II)] was monitored to evaluate the efficiency of photochemical energy conversion under actinic light. In contrast, non-photochemical quenching [Y(NPQ)] was analyzed to estimate the regulated thermal energy dissipation during dynamic light responses. The plants were dark-adapted overnight prior to measurements. Saturating pulses were applied for 720 ms at an intensity of 10, with a light intensity of 2, a modulation frequency of 1 Hz, a gain of 1, and a damping of 2.

### Determination of Nitrogen, Chlorophyll, Rubisco, Cl^-^, and NO_3_^+^

Leaf discs were excised from a fully expanded leaf of 7–8-week-old plants using a cork borer, immediately frozen in liquid nitrogen, and stored at −80 °C until analysis. The chlorophyll and RuBisCO contents per unit leaf area were quantified according to previously established protocols^40,41^. For nitrogen content determination, three leaves were oven-dried at 80 °C for 3 days, ground to a fine powder, and analyzed for carbon and nitrogen content using an elemental analyzer (SUMIGRAPH NC-22F; Sumika Chemical Analysis Service, Japan), as described previously^42^. For chloride (Cl□) determination, dried plant samples were ground to a fine powder. An aliquot of 80 mg (dry weight) of each sample was incubated in 1 mL of 0.5 N HNO□ in a 1.5 mL microtube at 30 °C for 24 h. After centrifugation, the supernatant was filtered and diluted tenfold. The chloride concentration was measured using an ICP–OES system (SPS3500, SII NanoTechnology, Japan) and quantified using the calibration curve method, according to the manufacturer’s instructions. For nitrate (NO□□) determination, plants were grown under 2 mM NO□□ hydroponic conditions or in soil. Nitrate was extracted from the roots and shoots using hot-water extraction, following a published protocol^43^, and subsequently analyzed by capillary electrophoresis using an Agilent 1600 CE system (Agilent Technologies, Santa Clara, CA, USA). Measurements were performed using an Agilent CE Inorganic Anions Solutions Kit (5063-6511) according to the manufacturer’s instructions.

### Transport assay

Y2H-based ABA transport assays were conducted essentially as described previously^44^, except that yeast growth was determined by monitoring the OD_600_ in liquid culture. Each transformant was precultured overnight at 30 ℃ in 96-well plates containing 100 μL of SD-LWU medium without shaking. After adding 100 μL of SD-LWU medium, the cells were resuspended by pipetting, and OD600 was adjusted to 0.1 with SD-LWUH medium containing 1 mM 3AT. An aliquot (20 μL) of the adjusted culture was then transferred to another 96-well plates containing 180 μL of SD-LWUH medium supplemented with 1 mM 3AT and 55.6 nM (+)-ABA (Tokyo Chemical Industry Co, Tokyo, Japan) (final concentration, 50 nM). The plates were incubated without shaking at 30 □, and OD600 was measured after 1, 2, and 3 days. OD600 was measured using a Varioskan LUX microplate reader (Thermo Fisher Scientific, Waltham, MA, USA).

For the direct measurement of ABA in yeast cells, *NPF2.4* and *NPF2.5* CDSs^44^ were cloned into pYES-DEST52, and the plasmid was transformed into *Saccharomyces cerevisiae* strain INVSc1 (Thermo Fisher, Waltham, MA, USA). Transport assays were conducted as described previously^26^. (+)-ABA (Tokyo Chemical Industry Co., Tokyo, Japan) was used as the substrate.

### ABA measurements

The hypocotyls of 7–8-week-old plants were cut with a razor blade, and the detached shoots were inserted into 15 mL tubes. The tubes were centrifuged at 6800 × *g* for 3 min at 5 °C, and the liquid at the bottom of the tube was collected as xylem sap. The collected xylem sap was then frozen at −80 °C until the analysis. Xylem sap (10 μL) was mixed with 100 μL of d_6_-ABA (5 pg/μL in 50% (v/v) MeOH) (OlchemIm, Czech Republic) and 90 μL of MeOH, and filtered with Cosmo Spin Filter G (Nacalai Tesque, Japan) that had been pretreated with 200 μL of MeCN and then with 200 μL of 80% (v/v) MeCN. The filter was washed with 100 μL of 80% (v/v) MeCN and combined with 200 μL of the flow-through obtained in the previous step to yield a xylem sap sample. The sample was dried in vacuo and dissolved in 10 μL of water for LC-MS/MS analysis with a mass spectrometer (Q Exactive; ThermoFisher Scientific, USA) coupled with an UHPLC (Vanquish; ThermoFisher Scientific, USA). The samples were separated on an ACQUITY PREMIER HSS T3 column (1.8µm VanGuardFIT 2.1×100 mm; Waters, MA, USA) with a gradient of solvent B (0.01% formic acid in MeCN) in combination with solvent A (0.06% acetic acid) indicated in Table S1. The flow rates of the LC are listed in Table S2. The settings of the mass spectrometer were as follows: mass tolerance (ppm) = 5, ion spray voltage (kV) = 2.5, collision energy (eV) = 10, S-lens radio frequency level = 50, resolution = 70,000, auto gain control target = 2×10^5^, maximum injection time (ms) = 200, isolation window (*m*/*z*) = 0.4, and ion mode = negative. Data were processed with Xcalibur 4.4 (ThermoFisher Scientific, USA). d_6_-ABA was identified based on transitions at *m*/*z* 269.2 156.108 and 159.130. Endogenous ABA was identified based on the *m*/*z* 263.1, 151.077, and 153.092 transitions. The retention times for d_6_-ABA and ABA were 13.1 min. The ABA levels were calculated based on the ratio of the internal standard (d_6_-ABA) to non-labeled ABA.

### GUS staining

After surface sterilization with 70% (v/v) ethanol and 5% (v/v) NaClO (0.25% active chlorine) containing 1% (w/v) Tween 20, the T_2_ seeds of *pNPF2.4:GUS* and *pNPF2.5:GUS* lines were sown on plates containing half-strength MS media, 0.8% (w/v) agar, and 1% (w/v) sucrose. The plates were incubated at 4 ℃ for 4 days in the dark and then incubated at 22 □ under a 10-h light/14-h dark photoperiod with a photosynthetic photon flux density of 50 μmol m^−2^ s^−1^. GUS staining was performed using a solution comprising 50 mM sodium phosphate buffer (pH 7.2), 10 mM EDTA, 0.05% (v/v) Triton X-100, 0.5 mM potassium ferrocyanide, 0.5 mM potassium ferricyanide, and 1 mM X-Gluc. Staining patterns were observed under a Leica M2-5 stereo microscope (Leica Camera AG, Wetzlar, Germany).

### Observation of GFP signals

After surface sterilization with 70% (v/v) ethanol and 5% (v/v) NaClO (0.25% active chlorine) containing 1% (w/v) Tween 20, T_2_ seeds of *pNPF2.4:gDNA-GFP* and *pNPF2.5:gDNA-GFP* lines were sown on plates containing half-strength MS medium, 1.2% (w/v) agar, and 1% (w/v) sucrose. The plates were incubated at 4 ℃ for 4 days in the dark and then incubated at 22 □ under an 8-h light/16-h dark photoperiod with a photosynthetic photon flux density of 50 μmol m^−2^ s^−1^. GFP fluorescence images were obtained using a confocal laser-scanning microscope SP8 (Leica, Wetzlar, Germany). GFP was excited at 488 nm, and emission signals were detected between 500 and 550 nm. For propidium iodide (PI) staining, roots were detached from seedlings and stained in a 30 µM PI solution. PI was excited at 488 nm, and emission signals were detected between 600 and 650 nm.

### Statistical Analysis

Statistical analysis was performed using R software (v.4.2.2; https://cran.r-project.org/bin/windows/base/) and GraphPad PRISM Software (v.11.0.0; GraphPad Software, San Diego, CA, USA). Data were analyzed using one-way ANOVA, followed by the Tukey–Kramer HSD test (*P* < 0.05) or Dunnett’s test (*P* < 0.05).

## Acknowledgments

Dr. Yoshihiro Ohmori (The University of Tokyo) for the quantification of Cl□ and NO□□. Dr. Tsuyoshi Nakagawa provided pGWB3 vector.

## Data Availability

All data are included in the manuscript and/or supporting information.

## References

1. Zeevaart, J. A. D. & Creelman, R. A. Metabolism and physiology of abscisic acid. Annu. Rev. Plant Physiol. Plant Mol. Biol. 39, 439–473 (1988). 10.1146/annurev.pp.39.060188.002255.

2. Shimadzu, S., Seo, M., Terashima, I. & Wataru, Y. Whole irradiated plant leaves showed faster photosynthesis induction than individually irradiated leaves via improved stomatal opening. Front. Plant Sci. 10, 1512 (2019). 10.3389/fpls.2019.01512

3. Farquhar, G. D. & Sharkey, T. D. Stomatal conductance and photosynthesis. Annu. Rev. Plant Physiol. 33, 317–345 (1982). 10.1146/annurev.pp.33.060182.001533.

4. Lawson, T. & Blatt, M. R. Stomatal size, speed, and responsiveness impact on photosynthesis and water use efficiency. Plant Physiol. 164, 1556–1570 (2014). 10.1104/pp.114.237107.

5. Pantin, F., Monnet, F., Jannaud, D., et al. The dual effect of abscisic acid on stomatal conductance and photosynthesis. Plant Physiol. 162, 1516–1529 (2013). 10.1111/nph.12013 Digital Object Identifier (DOI)

6. McAdam, S. A. M. & Brodribb, T. J. Linking turgor with ABA biosynthesis: implications for stomatal responses to vapor pressure deficit across land plants. Plant Physiol. 171, 2008–2016 (2016). 10.1104/pp.16.00380.

7. Seo, M. & Marion-Poll, A. Abscisic acid metabolism and transport in Abscisic Acid in Plants (eds Seo, M. & Marion-Poll, A.) (Elsevier, 2019) 1–49. 10.1016/bs.abr.2019.04.004

8. Kaiser, W. M. & Hartung, W. Uptake and release of abscisic acid by isolated photoautotrophic mesophyll cells, depending on pH gradients. Plant Physiol. 68, 202–206 (1981). 10.1104/pp.68.1.202.

9. Wilkinson, S. & Davies, W. J. Xylem sap pH increase: A drought signal received at the apoplastic face of the guard cell that involves the suppression of saturable abscisic acid uptake by the epidermal symplast. Plant Physiol. 113, 559–573 (1997). 10.1104/pp.113.2.559.

10. Kuromori, T., Seo, M. & Shinozaki, K. ABA transport and plant water stress responses. Trends Plant Sci. 21, 513–522 (2016). 10.1016/j.tplants.2018.04.001

11. Zhang, Y., Berman, A. & Shani, E. Plant hormone transport and localization: signaling molecules on the move. Annu. Rev. Plant Biol. 74, 453–479 (2023). 10.1146/annurev-arplant-070722-015329.

12. Davies, W. J., Kudoyarova, G. & Hartung, W. Long-distance ABA signaling and its relation to other signaling pathways in the detection of soil drying and the mediation of the plant’s response to drought. J. Plant Growth Regul. 24, 285–295 (2005). 10.1007/s00344-005-0103-1.

13. Dodd, I. C. Root-to-shoot signalling: assessing the roles of “up” in the up and down world of long-distance signalling in plants. Plant Soil 274, 251–270 (2005). 10.1007/s11104-004-0966-0.

14. Holbrook, N. M., Shashidhar, V. R., James, R. A. & Munns, R. Stomatal control in tomato with ABA-deficient roots: response of grafted plants to soil drying. J. Exp. Bot. 53, 1503–1514 (2002). 10.1093/jxb/53.373.1503.

15. Christmann, A., Weiler, E. W., Steudle, E. & Grill, E. A hydraulic signal in root-to-shoot signalling of water shortage. Plant J. 52, 167–174 (2007). 10.1111/j.1365-313X.2007.03234.x.

16. Manzi, M. et al. Root ABA accumulation in long-term water-stressed plants is sustained by hormone transport from aerial organs. Plant Cell Physiol. 56, 2457–2466 (2015). 10.1093/pcp/pcv161.

17. McAdam, S. A. M., Brodribb, T. J. & Ross, J. J. Shoot-derived abscisic acid promotes root growth. Plant Cell Environ. 39, 652–659 (2016). 10.1111/pce.12669.

18. Koiwai, H. et al. Tissue-specific localization of an abscisic acid biosynthetic enzyme, AAO3, in Arabidopsis. Plant Physiol. 134, 1697–1707 (2004). 10.1104/pp.103.036970.

19. Endo, A. et al. Drought induction of Arabidopsis 9-cis-epoxycarotenoid dioxygenase occurs in vascular parenchyma cells. Plant Physiol. 147, 1984–1993 (2008). 10.1104/pp.108.116632.

20. Kuromori, T., Sugimoto, E. & Shinozaki, K. Intertissue signal transfer of abscisic acid from vascular cells to guard cells. Plant Physiol. 164, 1587–1592 (2014). 10.1104/pp.114.235556.

21. Merilo, E. et al. Stomatal VPD response: there is more to the story than ABA. Plant Physiol. 176, 851–864 (2018). 10.1104/pp.17.00912.

22. Bauer, H. et al. The stomatal response to reduced relative humidity requires guard cell-autonomous ABA synthesis. Curr. Biol. 23, 53–57 (2013). 10.1016/j.cub.2012.11.022.

23. Kuromori, T., et al. ABC transporter AtABCG25 is involved in abscisic acid transport and responses. Proc. Natl Acad. Sci. U. S. A. 107, 2361–2366 (2010). 10.1073/pnas.0912516107.

24. Kang, J., et al. PDR-type ABC transporter mediates cellular uptake of the phytohormone abscisic acid. Proc. Natl Acad. Sci. U. S. A. 107, 2355–2360 (2010). 10.1073/pnas.0909222107.

25. Kanno, Y., et al. Identification of an abscisic acid transporter by functional screening using the receptor complex as a sensor. Proc. Natl Acad. Sci. U. S. A. 109, 9653–9658 (2012). 10.1073/pnas.1203567109.

26. Shimizu, T., Kanno, Y., Suzuki, H., Watanabe, S. & Seo, M. Arabidopsis NPF4.6 and NPF5.1 control leaf stomatal aperture by regulating abscisic acid transport. Genes 12, 885 (2021). 10.3390/genes12060885.

27. Corratgé-Faillie, C. & Lacombe, B. Substrate (un)specificity of Arabidopsis NRT1/PTR FAMILY (NPF) proteins. J. Exp. Bot. 68, 3107–3113 (2017). 10.1093/jxb/erw499.

28. Li, B. et al. Identification of a stelar-localized transport protein that facilitates root-to-shoot transfer of chloride in Arabidopsis. Plant Physiol. 170, 1014–1029 (2016). 10.1104/pp.15.01163.

29. Li, B. et al. AtNPF2.5 modulates chloride (Cl−) efflux from roots of Arabidopsis thaliana. Front. Plant Sci. 7, 2013 (2017). 10.3389/fpls.2016.02013.

30. Binenbaum, J. et al. Gibberellin and abscisic acid transporters facilitate endodermal suberin formation in Arabidopsis. Nat. Plants 9, 785–802 (2023). 10.1038/s41477-023-01391-3.

31. Tsay, Y.-F., Chiu, C.-C., Tsai, C.-B., Ho, C.-H. & Hsu, P.-K. Nitrate transporters and peptide transporters. FEBS Lett. 581, 2290–2300 (2007). 10.1016/j.febslet.2007.04.047.

32. Takahashi, F. et al. A small peptide modulates stomatal control via abscisic acid in long-distance signalling. Nature 556, 235–238 (2018). 10.1038/s41586-018-0009-2.

33. Sun, S.-K. et al. The plastid cysteine synthase complex regulates ABA biosynthesis and stomatal closure in Arabidopsis. Nat. Commun. 16, 8960 (2025). 10.1038/s41467-025-64705-3.

34. Zhang, Y. et al. ABA homeostasis and long-distance translocation are redundantly regulated by ABCG ABA importers. Sci. Adv. 7, eabf6069 (2021). 10.1126/sciadv.abf6069.

35. Yang, Q. et al. Abscisic acid root-to-shoot translocation by transporter AtABCG25 mediates stomatal movements in Arabidopsis. Plant Physiol. 195, 671–684 (2024). 10.1093/plphys/kiae073.

36. Mehra, P. et al. Hydraulic flux-responsive hormone redistribution determines root branching. Science 378, 762–768 (2023). 10.1126/science.add3771

37. Winter, D. et al. An “Electronic Fluorescent Pictograph” browser for exploring and analyzing large-scale biological data sets. PLOS One 2, e718 (2007). 10.1371/journal.pone.0000718.

38. Nakagawa, T. et al. Development of series of gateway binary vectors, pGWBs, for realizing efficient construction of fusion genes for plant transformation. J. Biosci. Bioeng. 104, 34–41 (2007). 10.1263/jbb.104.34.

39. Shimada, T. L., Shimada, T. & Hara-Nishimura, I. A rapid and non-destructive screenable marker, FAST, for identifying transformed seeds of Arabidopsis thaliana. Plant J. 61, 519–528 (2010). 10.1111/j.1365-313X.2009.04060.x.

40. Porra, R. J., Thompson, W. A. & Kriedemann, P. E. Determination of accurate extinction coefficients and Simultaneous equations for assaying chlorophylls a and b extracted with four different solvents: verification of the concentration of chlorophyll standards by atomic absorption spectroscopy. Biochim. Biophys. Acta Bioenerg. 975, 384–394 (1989). 10.1016/S0005-2728(89)80347-0.

41. Kimura, H., Hashimoto-Sugimoto, M., Iba, K., Terashima, I. & Yamori, W. Improved stomatal opening enhances photosynthetic rate and biomass production in fluctuating light. J. Exp. Bot. 71, 2339–2350 (2020). 10.1093/jxb/eraa090.

42. Katsuhama, N. et al. PROTON ATPASE TRANSLOCATION CONTROL 1-mediated H + -ATPase translocation boosts plant growth under drought by optimizing root and leaf functions. PNAS Nexus 4, pgaf151 (2025). 10.1093/pnasnexus/pgaf151.

43. Hachiya, T. & Okamoto, Y. Simple spectroscopic determination of nitrate, nitrite, and ammonium in Arabidopsis thaliana. Bio-Protoc. 7, e2280 (2017). 10.21769/BioProtoc.2280.

44. Chiba, Y. et al. Identification of Arabidopsis thaliana NRT1/PTR FAMILY (NPF) proteins capable of transporting plant hormones. J. Plant Res. 128, 679–686 (2015). 10.1007/s10265-015-0710-2.

