## Supplementary table and figures for "Root-to-shoot ABA transport by NPF2.4 and NPF2.5 links stomatal regulation to photosynthetic performance"

Supplemental Table 1. Percent of solvent B during the LC-MS/MS analysis.

| Time (min) | 0 | 10 | 11 | 15 | 16 | 16.1 | 16.999 | 17 | 21 |
| --- | --- | --- | --- | --- | --- | --- | --- | --- | --- |
| % B | 5 | 25 | 29.5 | 35.5 | 44.5 | 99 | 99 | 5 | 5 |

Supplemental Table 2. Flow rate of LC during the LC-MS/MS analysis.

| Time (min) | 0 | 16 | 16.1 | 16.9 | 16.999 | 21 |
| --- | --- | --- | --- | --- | --- | --- |
| Flow rate (mL/min) | 0.25 | 0.25 | 0.4 | 0.4 | 0.25 | 0.25 |

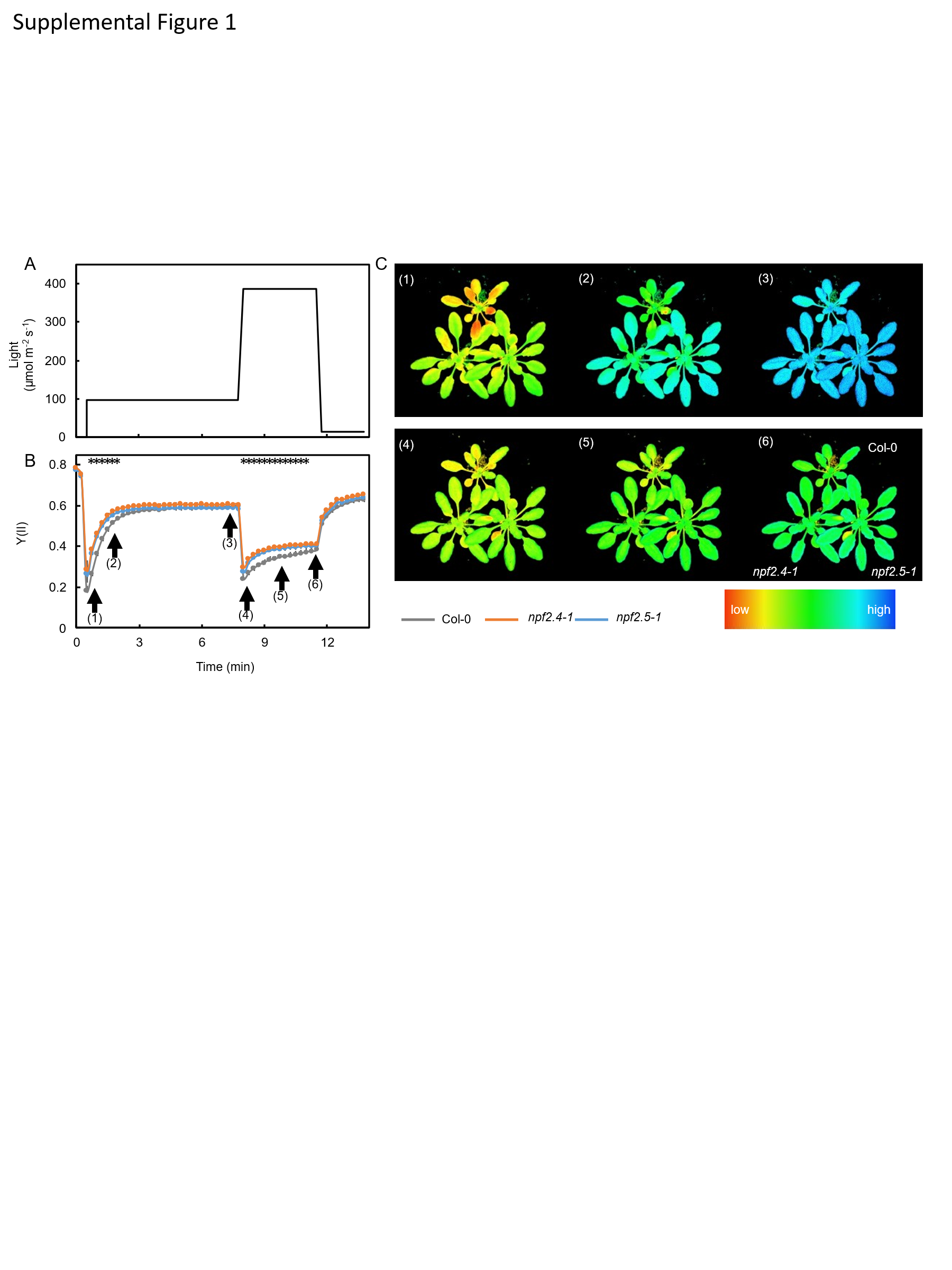

**Supplemental Figure 1. Photosynthetic responses to fluctuating light.**

(A) Light fluctuation regime used for chlorophyll fluorescence imaging.

(B) Effective quantum yield of photosystem II, Y(II), under the light fluctuation shown in (A).

(C) Representative chlorophyll fluorescence images corresponding to time points (1)–(6) indicated in (B).

Measurements were performed using plants grown for 30 days at 23 °C under a light intensity of 130 µmol m⁻² s⁻¹ with a 10-h photoperiod. Values represent mean ± SEM. Different letters indicate significant differences at *P* < 0.05 as determined by Tukey–Kramer HSD test (*n* = 6). When Col-0 shows a statistically significant difference compared with both *npf2.4* and *npf2.5*, this is indicated by an asterisk (*).

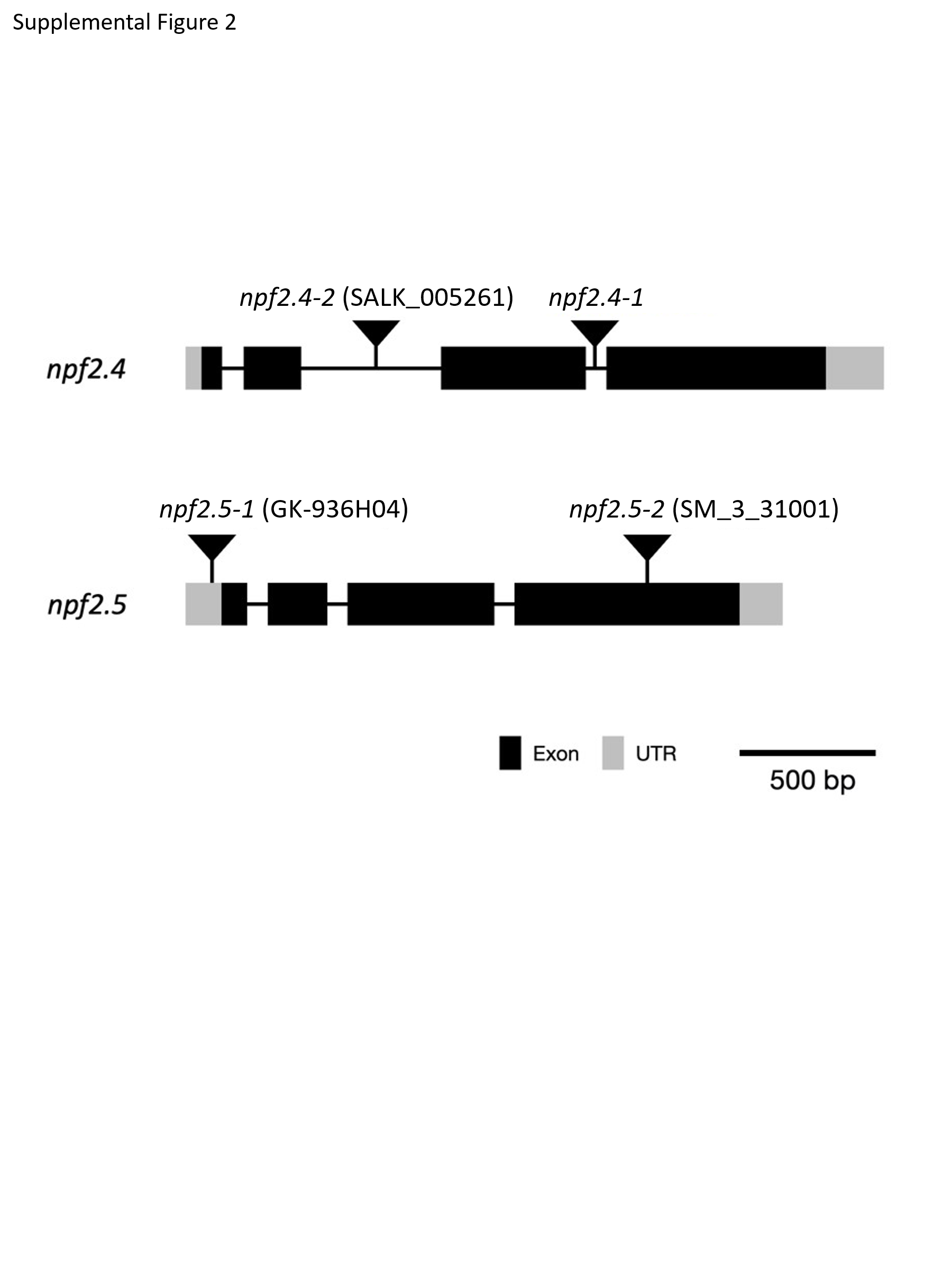

**Supplemental Figure S2. Structure of *NPF2.4* and *NPF2.5* genes and the positions of T-DNA or transposon insertion.**

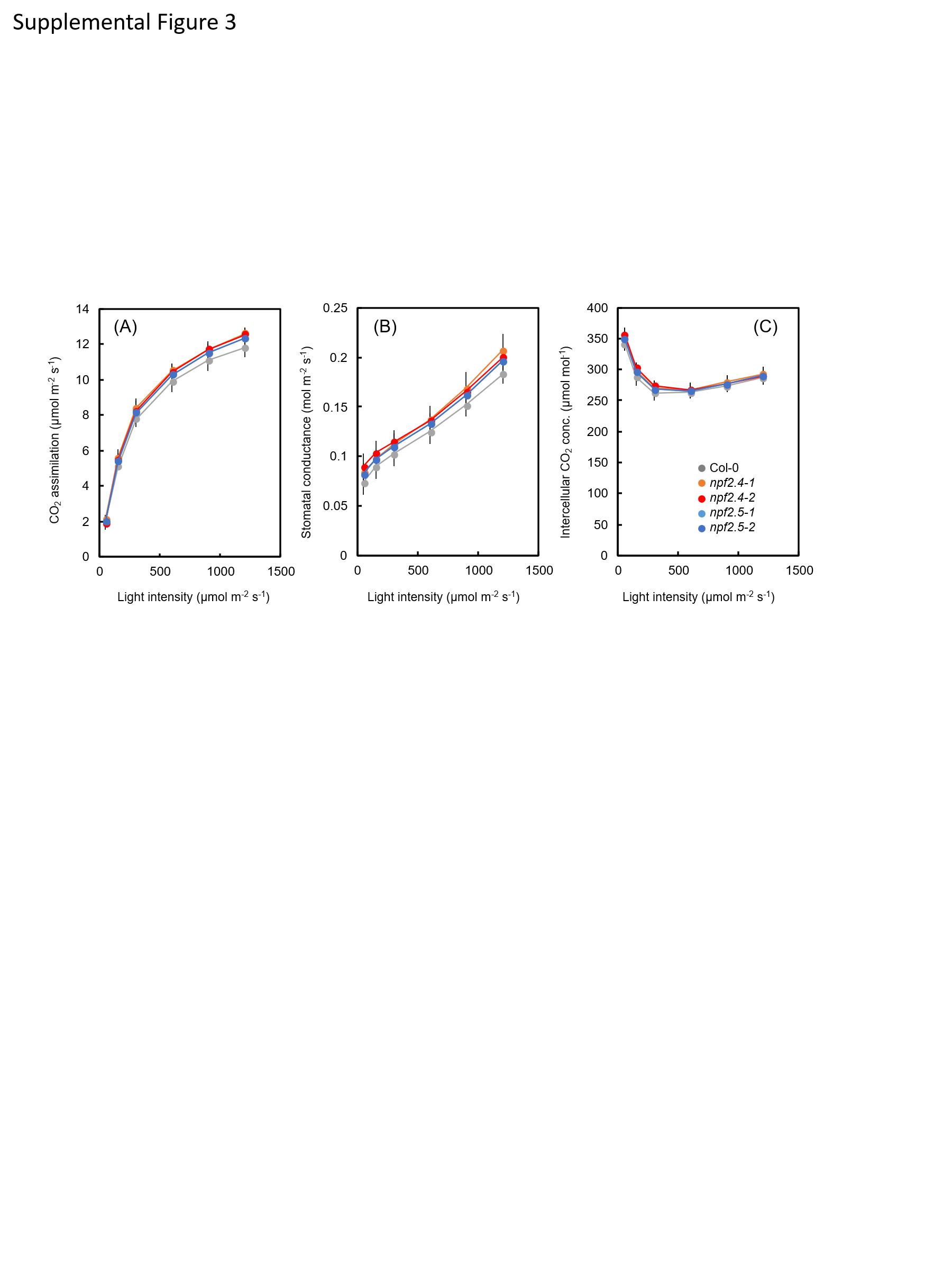

**Supplemental Figure 3. Light response of photosynthesis under steady-state conditions.**

(A) Light response of CO₂ assimilation rate.

(B) Light response of stomatal conductance.

(C) Light response of intercellular CO₂ concentration.

Values represent mean ± SEM. No significant differences were detected at any light intensity as determined by Tukey–Kramer HSD test (*n* = 6).

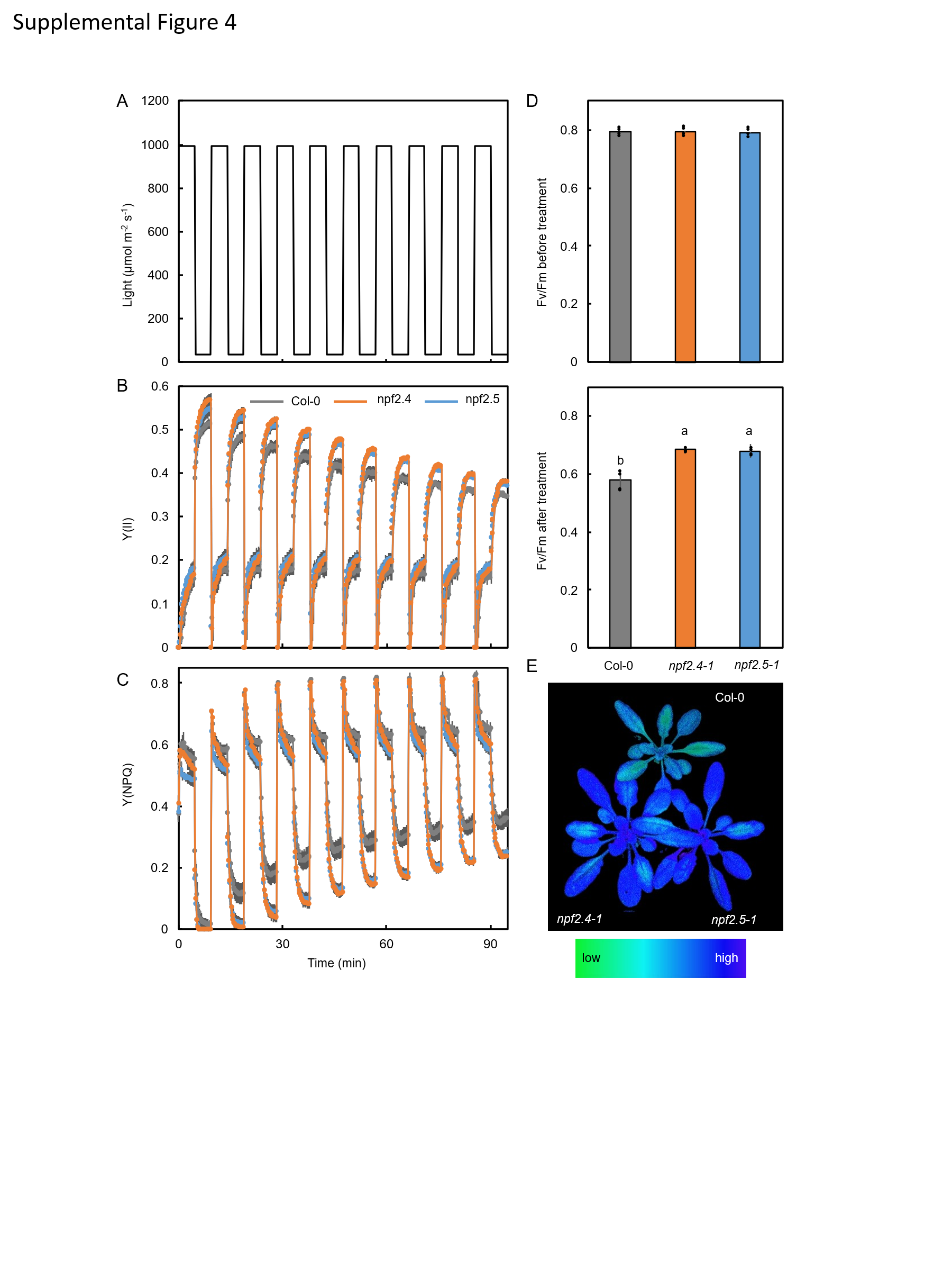

**Supplemental Figure 4. Photosynthetic responses to fluctuating light.**

(A) Light fluctuation regime used for chlorophyll fluorescence imaging.

(B) Effective quantum yield of photosystem II, Y(II), under the fluctuating light shown in (A).

(C) Quantum yield of regulated thermal dissipation in photosystem II, Y(NPQ), under the fluctuating light shown in (A).

(D) Maximum quantum yield of PSII (Fv/Fm) measured before and after fluctuating light stress shown in (A).

(E) Representative images of Fv/Fm after fluctuating light stress shown in (A).

Measurements were performed using plants grown for 30 days at 23 °C under a light intensity of 130 µmol m⁻² s⁻¹ with a 10-h photoperiod. Values represent mean ± SEM. Different letters indicate significant differences at *P* < 0.05 as determined by Tukey–Kramer HSD test (*n* = 6).

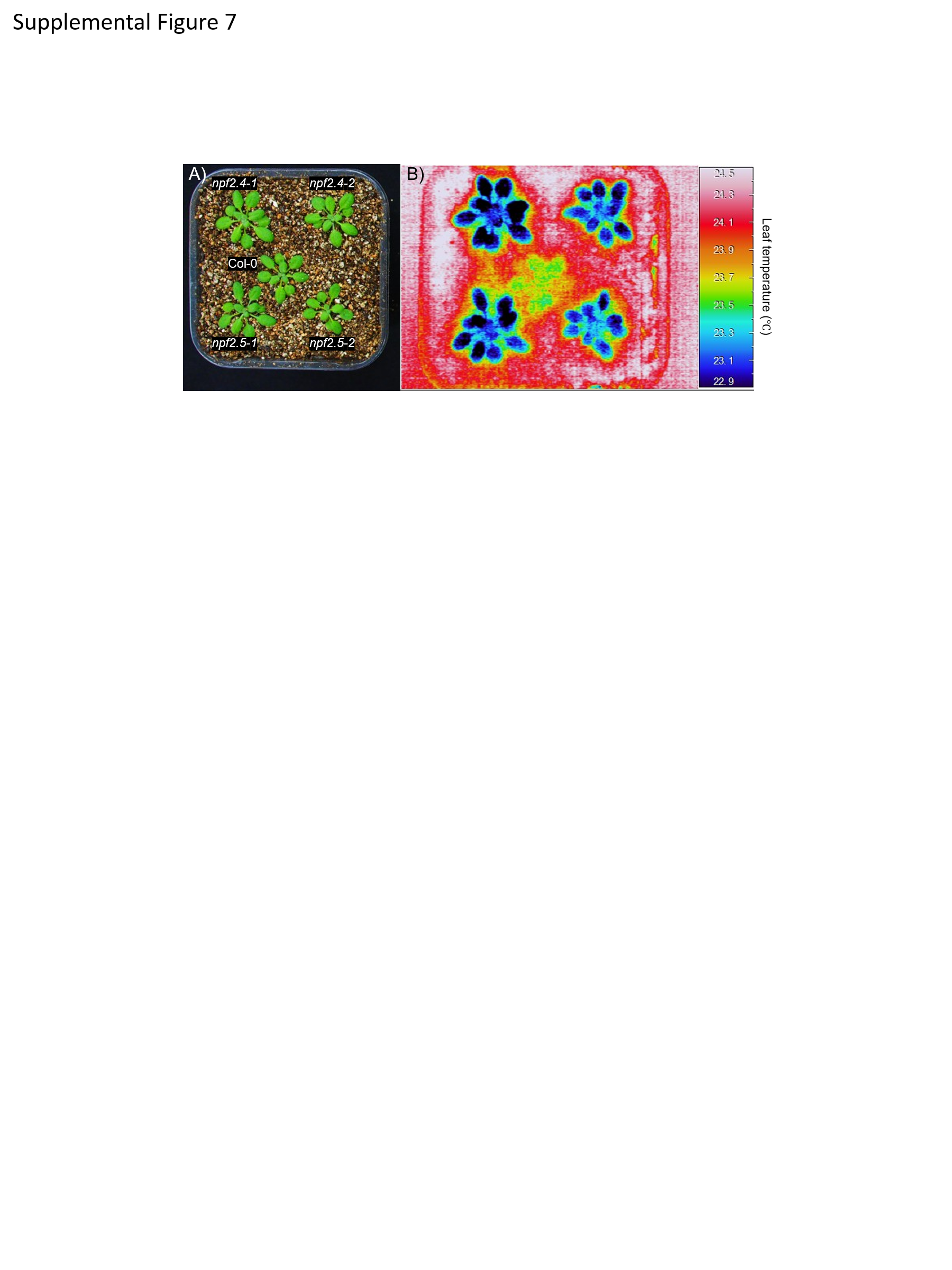

**Supplemental Figure 5. Plant growth and thermal imaging.**

(A) Representative images of plant growth and

(B) corresponding thermal images obtained by infrared thermography of 25-day-old plants.

Thermal images were acquired 1 h after the onset of illumination under growth light conditions. The summarized thermal imaging and quantitative data are presented in Figure 2.

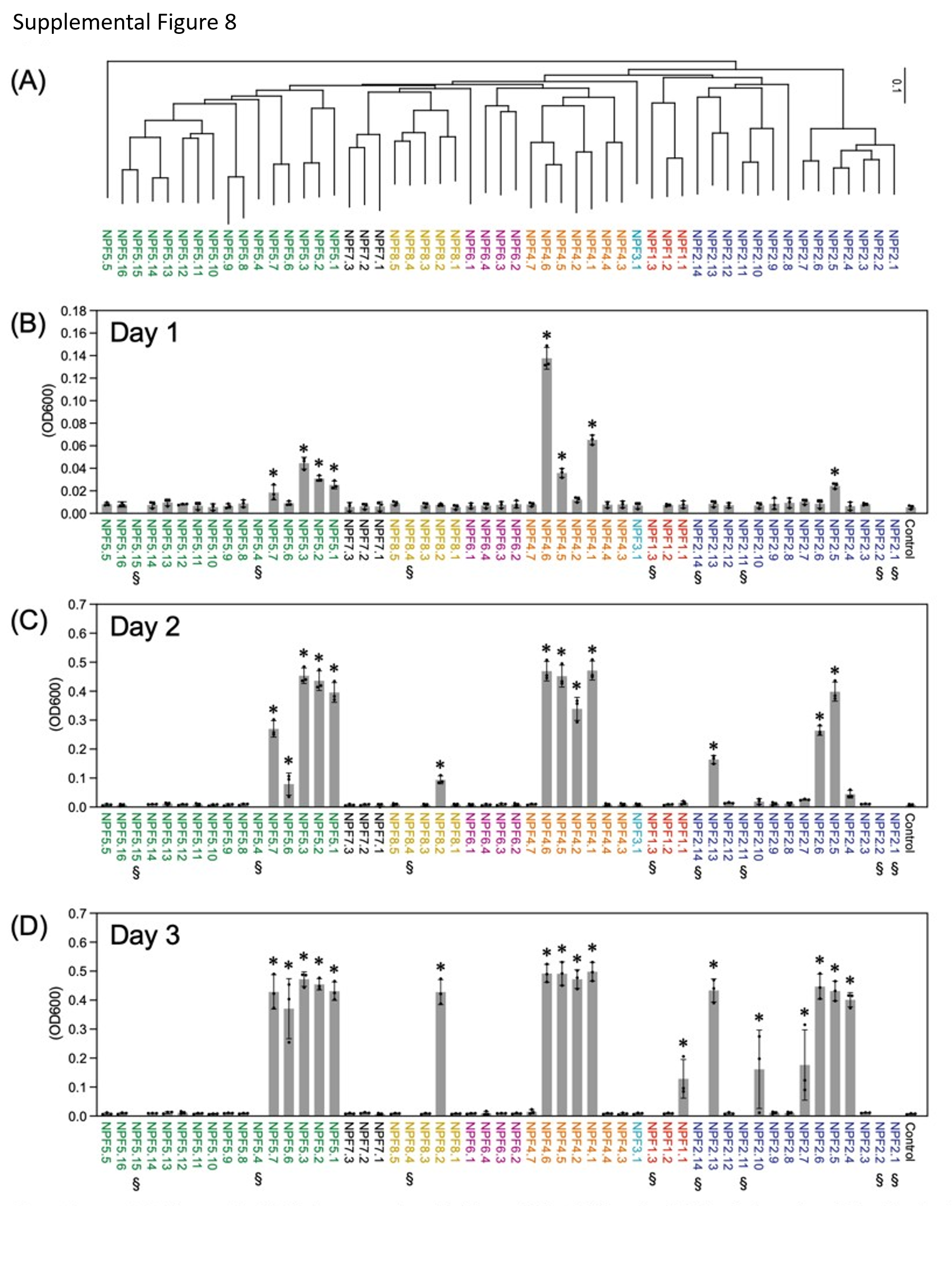

**Supplemental Figure 6. ABA transport activities of Arabidopsis NPFs determined by the indirect assay.**

(A) Phylogenetic tree of 53 Arabidopsis NPFs.

(B) - (D) Growth (OD_600_) of yeast strains expressing the Y2H-based ABA sensor and NPF proteins in the presence of 50 nM ABA in liquid culture. Control cells contain the empty vector instead of NPFs. Yeast growth was determined 1 (B), 2 (C) and 3 (D) days after inoculation. NPFs marked with section signs (§) were not analyzed. Asterisk indicates significant difference against Control (P<0.05) by Dunnett’s multiple comparison test (n = 3).

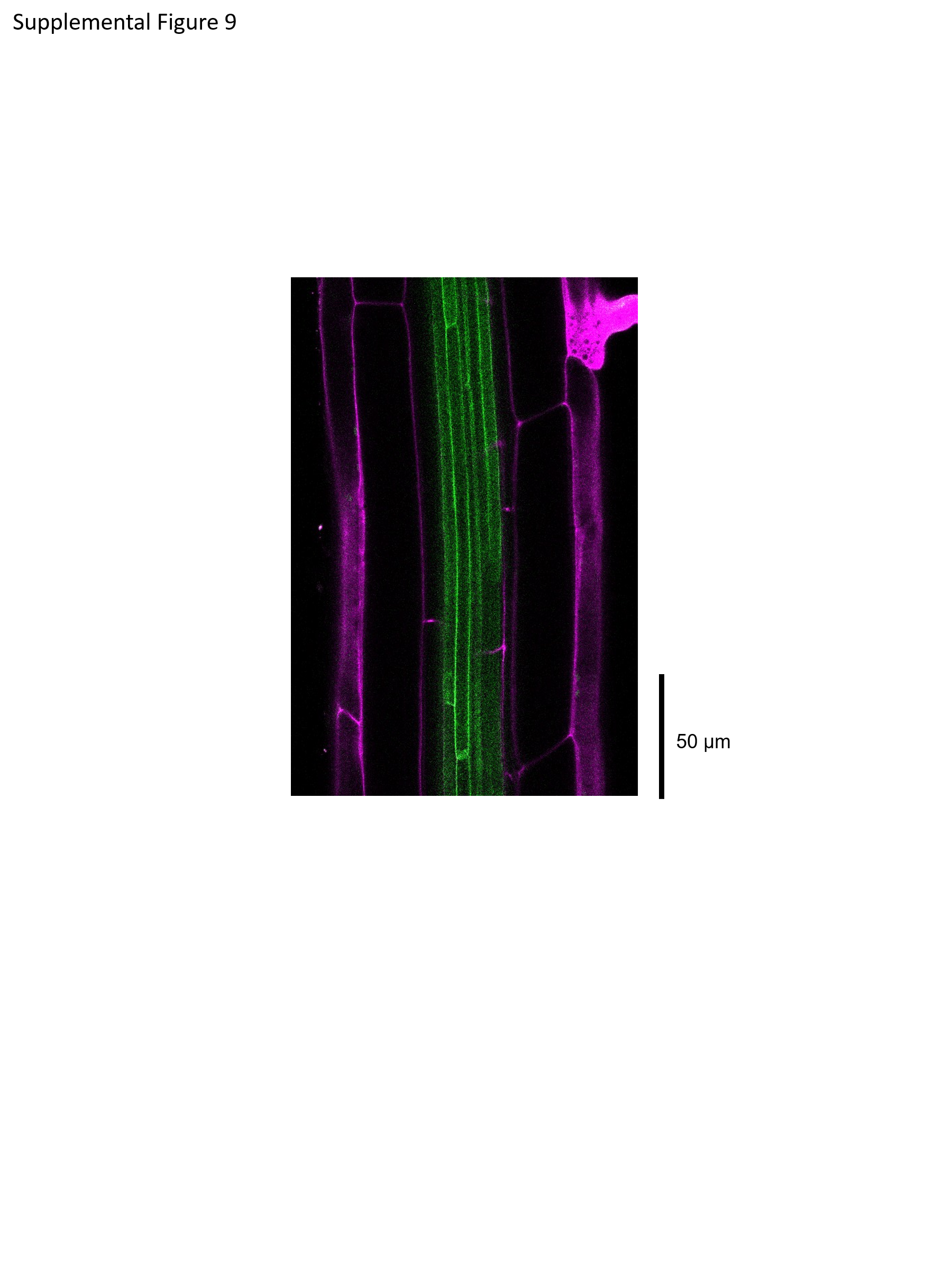

**Supplemental Figure 7. Plasma membrane localization of GFP signals in the pericycle cells of *pNPF2.4:gNPF2.4-GFP* plants.**

Green color indicates GFP signals detected in 10-day-old roots of *pNPF2.4:gNPF2.4-GFP* plants. Magenta color indicates Pi staining of cell walls.

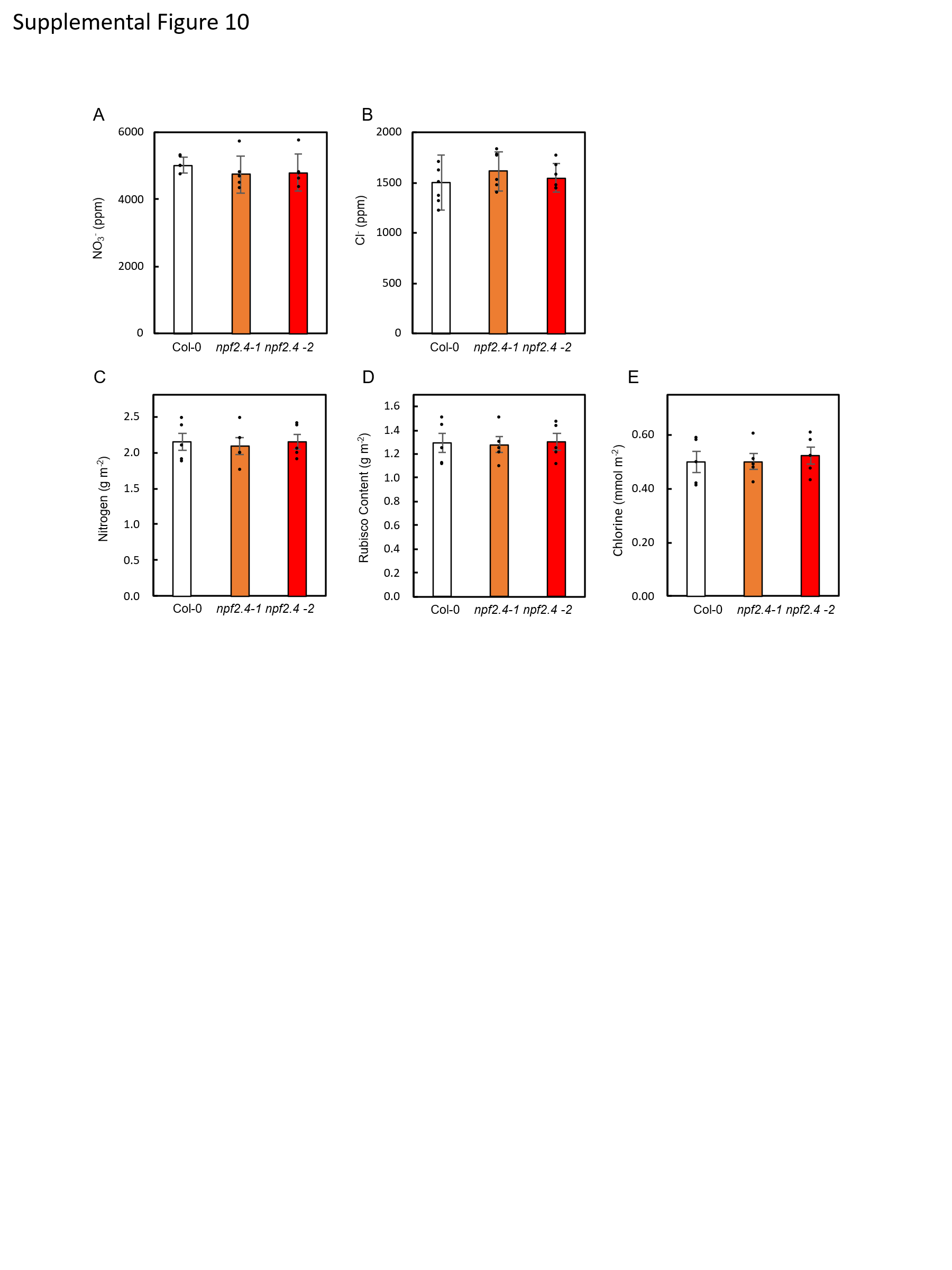

**Supplemental Figure 8. Nitrate and chloride contents in leaves.**

(A) Nitrate (NO₃⁻) content in leaves.

(B) Chloride (Cl⁻) content in leaves.

(C) Total leaf nitrogen content.

(D) Rubisco content.

(E) Chlorophyll content.

Measurements were performed using plants grown for 30 days at 23 °C under a light intensity of 130 µmol m⁻² s⁻¹ with a 10-h photoperiod. Values represent mean ± SEM. No significant differences were detected as determined by Tukey–Kramer HSD test (*n* = 6).

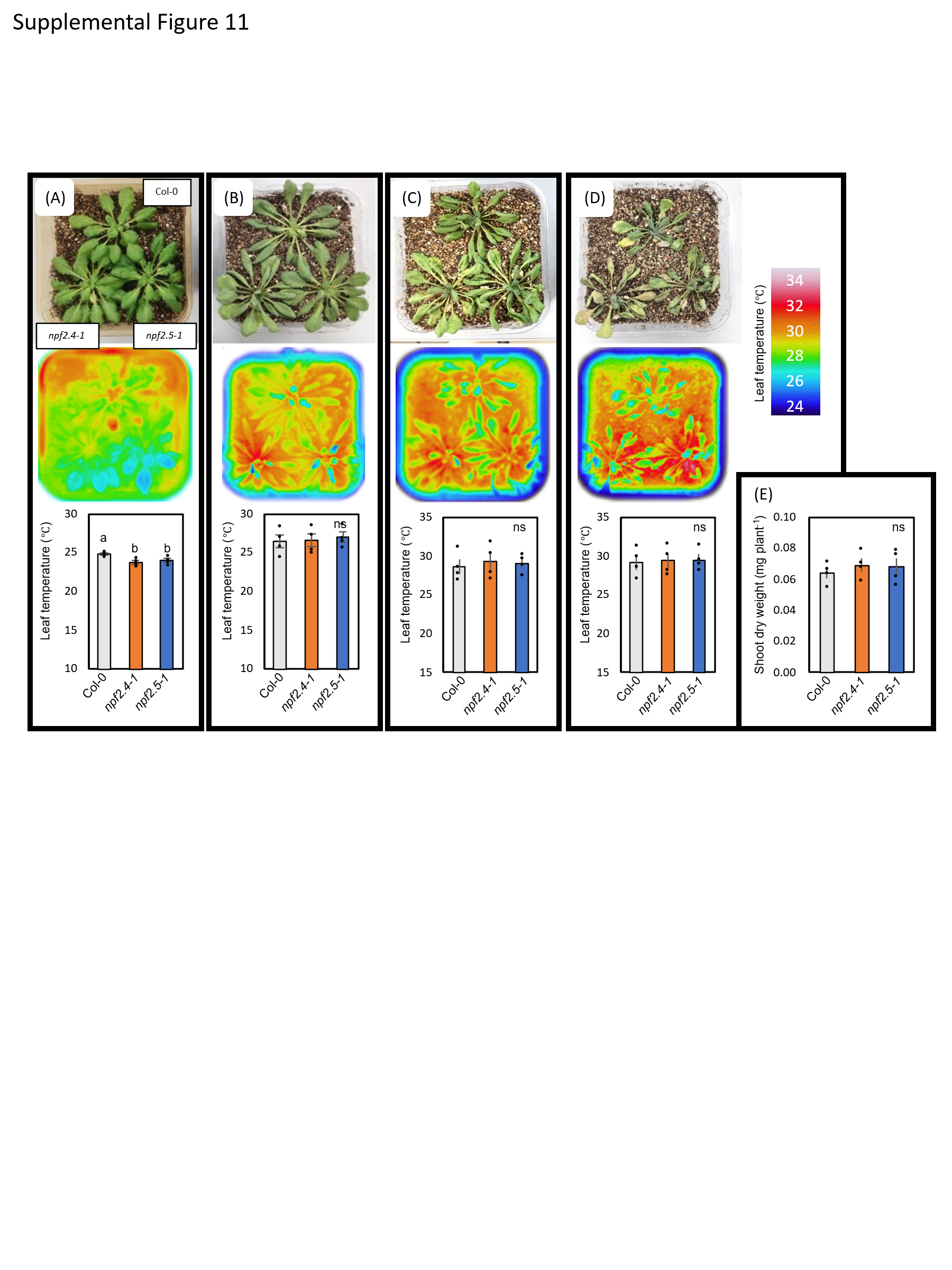

**Supplemental Figure 9. Resistance to long-term water deficit in npf2.4 and npf2.5 mutants**

(A) Representative images of plant growth (left), corresponding thermal images (middle), and quantified leaf temperatures (right) of Col-0, *npf2.4-1*, and *npf2.5-1* plants grown for 30 days under well-watered conditions. Plants were cultivated under controlled environmental conditions at 23 °C, 65% relative humidity, a 10-h light/14-h dark photoperiod, and a photosynthetic photon flux density of 130 μmol m⁻² s⁻¹, under ambient CO₂ concentration (400 μmol mol⁻¹).

(B) Representative images of plant growth, thermal images, and leaf temperature measurements after withholding water for 10 days starting from 20 days after sowing.

(C) Representative images of plant growth, thermal images, and leaf temperature measurements after 12 days of water withholding.

(D) Representative images of plant growth, thermal images, and leaf temperature measurements after 14 days of water withholding.

(E) Dry weight of Col-0, *npf2.4-1*, and *npf2.5-1* plants harvested at the stage shown in (D).

Values represent mean ± SEM (n = 4).

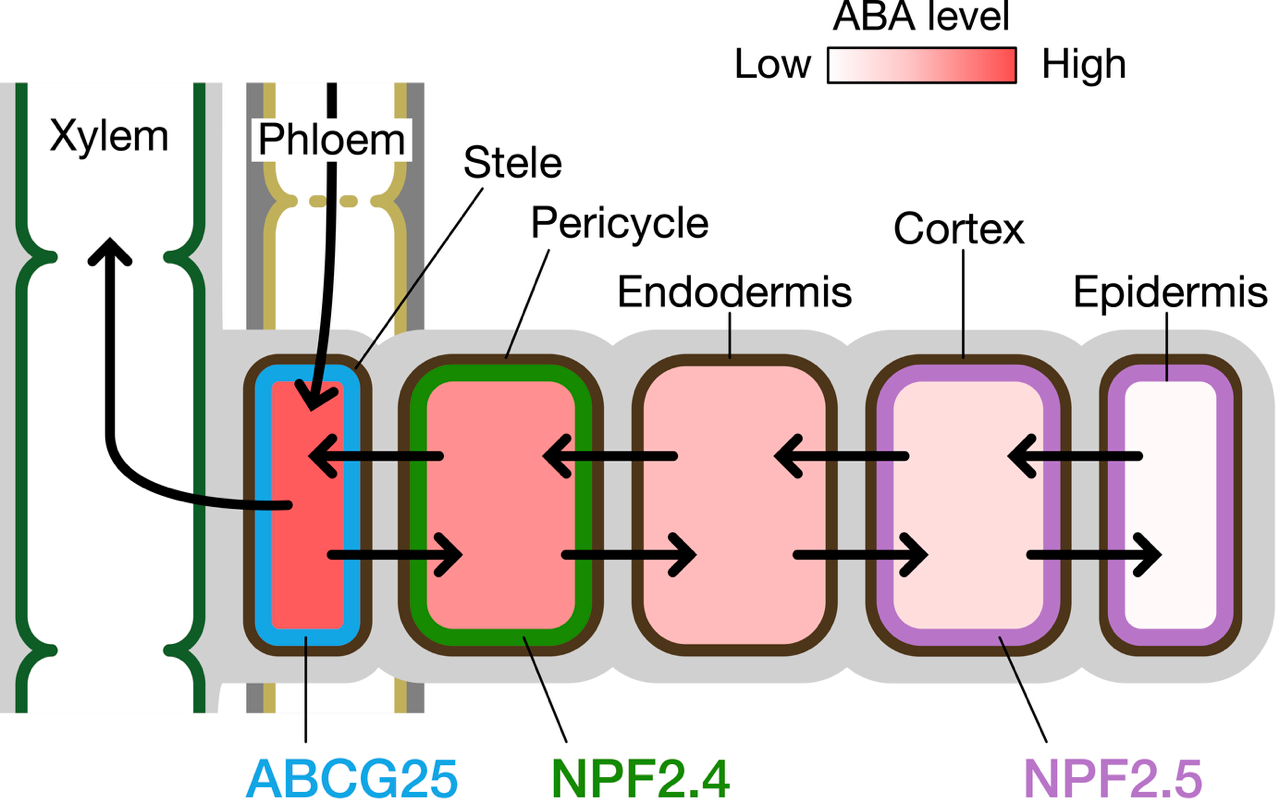

**Supplemental Figure 10. Possible movement of ABA and spatial expression patterns of NPF2.4 and NPF2.5 within root tissues.**

ABA is synthesized in root vascular tissues or translocated from shoots via phloem. ABA can be transported radially in both inward and outward directions in roots. NPF2.4 is predominantly expressed in pericycle cells, whereas NPF2.5 is expressed in cortex and epidermal cells. Both proteins mediate cellular ABA uptake.

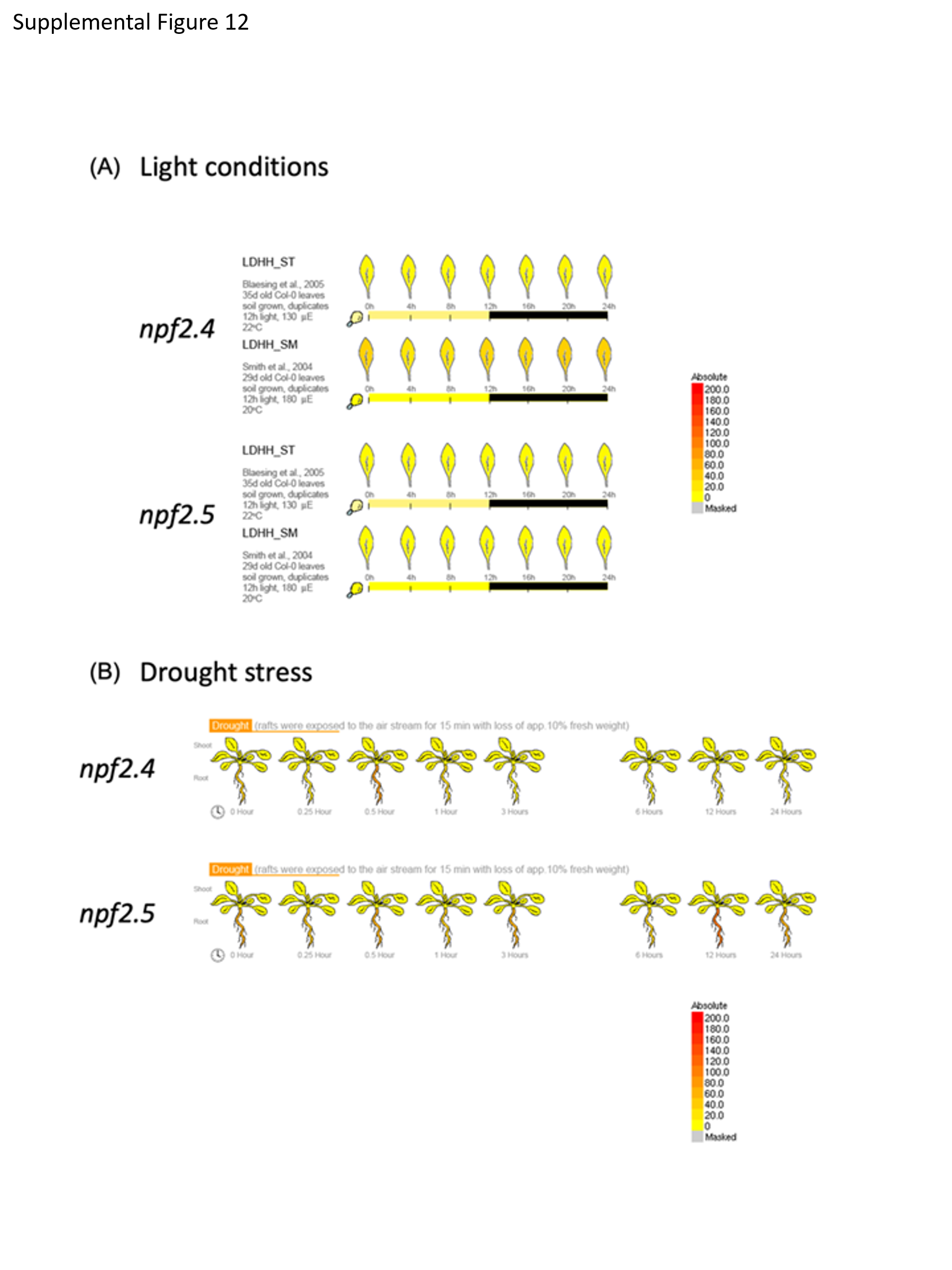

**Supplemental Figure 11.** Expression levels of *NPF2.4* and *NPF2.5*.

(A) Expression under a 12 h light/12 h dark photoperiod.

(B) Expression under drought stress conditions.

Images were obtained from Arabidopsis eFP Browser (https://bar.utoronto.ca/efp/cgi-bin/efpWeb.cgi).
